# SARS-CoV-2 Quasispecies provides insight into its genetic dynamics during infection

**DOI:** 10.1101/2020.08.20.258376

**Authors:** Fengming Sun, Xiuhua Wang, Shun Tan, Yunjie Dan, Yanqiu Lu, Juan Zhang, Junli Xu, Zhaoxia Tan, Xiaomei Xiang, Yi Zhou, Weiwei He, Xing Wan, Wei Zhang, Yaokai Chen, Wenting Tan, Guohong Deng

## Abstract

A novel coronavirus disease (COVID-19) caused by SARS-CoV-2 has been pandemic worldwide. The genetic dynamics of quasispecies afford RNA viruses a great fitness on cell tropism and host range. However, no quasispecies data of SARS-CoV-2 have been reported yet. To explore quasispecies haplotypes and its transmission characteristics, we carried out single-molecule real-time (SMRT) sequencing of the full-length of SARS-CoV-2 spike gene within 14 RNA samples from 2 infection clusters, covering first-to third-generation infected-patients. We observed a special quasispecies structure of SARS-CoV-2 (modeled as ‘One-King’): one dominant haplotype (mean abundance ~70.15%) followed by numerous minor haplotypes (mean abundance < 0.10%). We not only discovered a novel dominant haplotype of F^1040^ but also realized that minor quasispecies were also worthy of attention. Notably, some minor haplotypes (like F^1040^ and currently pandemic one G^614^) could potentially reveal adaptive and converse into the dominant one. However, minor haplotypes exhibited a high transmission bottleneck (~6% could be stably transmitted), and the new adaptive/dominant haplotypes were likely originated from genetic variations within a host rather than transmission. The evolutionary rate was estimated as 2.68-3.86 × 10^−3^ per site per year, which was larger than the estimation at consensus genome level. The ‘One-King’ model and conversion event expanded our understanding of the genetic dynamics of SARS-CoV-2, and explained the incomprehensible phenomenon at the consensus genome level, such as limited cumulative mutations and low evolutionary rate. Moreover, our findings suggested the epidemic strains may be multi-host origin and future traceability would face huge difficulties.

## Introduction

Coronaviruses are enveloped, single, and positive-stranded RNA viruses that can be further classified into four genera: *alphacoronaviruses*, *betacoronaviruses*, *gammacoronaviruses*, and *deltacoronaviruses*^1^. The *betacoronaviruses* have a zoonotic potential and can be transmitted from animals to humans causing a novel, severe respiratory disease^2^. The severe acute respiratory syndrome (SARS) caused by SARS-related coronavirus (SARS-CoV) was broken out in 2002, and globally resulted in nearly 8000 laboratory-confirmed cases and 800 deaths^3,4^. The natural reservoir of SARS-CoV was presumed to be *Rhinolophus Sinensis* (Coronavirus strains: RsSHC014 and Rs3367)^5^. In 2012, another *betacoronaviruse* (MERS-CoV) caused the outbreak of Middle East respiratory syndrome (MERS) ^6,7^. By November 2019, the MERS have resulted in 2494 cases and 858 deaths worldwide^8^. The origin of MERS-CoV remains elusive, but studies have shown that humans were infected by direct or indirect contact with the infected dromedary camels in Arabian Peninsula (WHO). Although the two large-scale epidemics have gradually subsided, the threat from zoonotic *betacoronaviruses* always exists due to its inherent characteristics (genetic diversity) and frequently contact between humans and animals (hunting and domestication).

In December 2019, Wuhan, China, has emerged unknown pneumonia cases with the main symptoms of fever, fatigue, cough, and poor breath (COVID-19). The epidemic has been confirmed to be caused by a novel *betacoronaviruses* called SARS-CoV-2 ^9^. Genome sequences of SARS-CoV-2 showed a high similarity (~0.96) with bat coronavirus RaTG13, indicating a potential bat origin of the virus^10,11.^ The COVID-19 is still pandemic, and there were over 15 million confirmed cases and 600,000 deaths worldwide by August, 2020. Although antiviral drugs and vaccines are under development^12,13^, the actual work will face many challenges like long-periodic clinical trials and the risk of poor effects caused by high mutation rate. To guide and obtain effective antiviral strategies, it is therefore crucial to study and understand the underlying genetic mechanisms of fitness during infection.

In the consensus genome level, SARS-CoV-2 has been reported with limited variations^14–17^. RNA virus population in a host does not consist of a consensus single haplotype, but rather an ensemble of related sequences termed quasispecies. The dynamic changes of quasispecies affords RNA viruses a greater probability to change their cell tropism or host range or to overcome internal or external selective constraints^18^. The process is likely achieved by changing the genetic characteristics of key functional genes, such as the spike glycoprotein gene^19^. A novel amino mutation D614G within spike protein has currently found to be widely spread in newly infected patients, and its infectious ability reveals 2.6-9.3 times stronger than the early ancestor D^614^ haplotype^20–22^. Spike gene could be primarily determined its strong infectious ability of SARS-CoV-2 through binding the receptor angiotensin-converting enzyme 2 (ACE2) of host cell^23,24^. Study the quasispecies can not only observe the dynamic fitness of SARS-CoV-2, but also widely discover the potential advantage mutations (like G^614^) in advance, which were crucial for virus prevention, such as drug design and vaccine development. However, no data have reported its quasispecies yet.

Therefore, the aim of the study is to investigate the quasispecies features of SARS-CoV-2. We applied PicBio single-molecule real-time (SMRT) technology and circular consensus sequences (CCS) algorithm to obtain quasispecies haplotypes. The included 9 cases and its related 14 samples were carefully prepared to achieve two major research objectives. Firstly, to fully trace the quasispecies features and its dynamics during transmission (between hosts), our cases were screened from two single-source infection sets (containing two familial clusters), and covered first-to third-generation patients. Secondly, to facilitate the discovery of quasispecies differences between sampling types (within host), RNAs were designed to extract from nasopharyngeal swabs (NS), sputum (SP), and stool (ST). Moreover, we have also predicted the potential advantage genotypes and the evolutionary rate at the quasispecies level. Our results provided insight into the genetic fitness of SARS-CoV-2 and also provided important information for the follow-up prevention.

## Results

### Single-molecule real-time sequencing of the SARS-CoV-2 spike gene

For all 14 samples (Fig. 1 and **Supplementary Table1**), the SMRT platform produced 26.97 Gb subreads (Mean: 1.93±0.88 Gb) and the corresponding 379,079 CCS (5,044 ~ 52,739) (**Supplementary Table 2**). Metagenomic sequencing revealed that more than 98% of quasispecies RNA contains the identical designed primer sequences, indicating the amplified segments (subreads and CCS) have no obvious primer bias (**Supplementary Fig. 1**). Through increasing the pass number, the main type of sequencing errors within subreads (gaps: insertions and deletions) could be self-corrected during the generation of CCS. Statistics showed that the average gap number of CCS decreases to a low stable value (median: 23, account for 0.60% of full length) when the pass number equaled 5 (Fig. 2a). Therefore, only CCS with pass ≥5 were considered for further analysis, which accounted for 74.83% (283,655) of the total number and the related subreads accounted for 90.54% of the total sequencing data (Fig. 2b and **Supplementary Table 2**).

**Figure 1.**
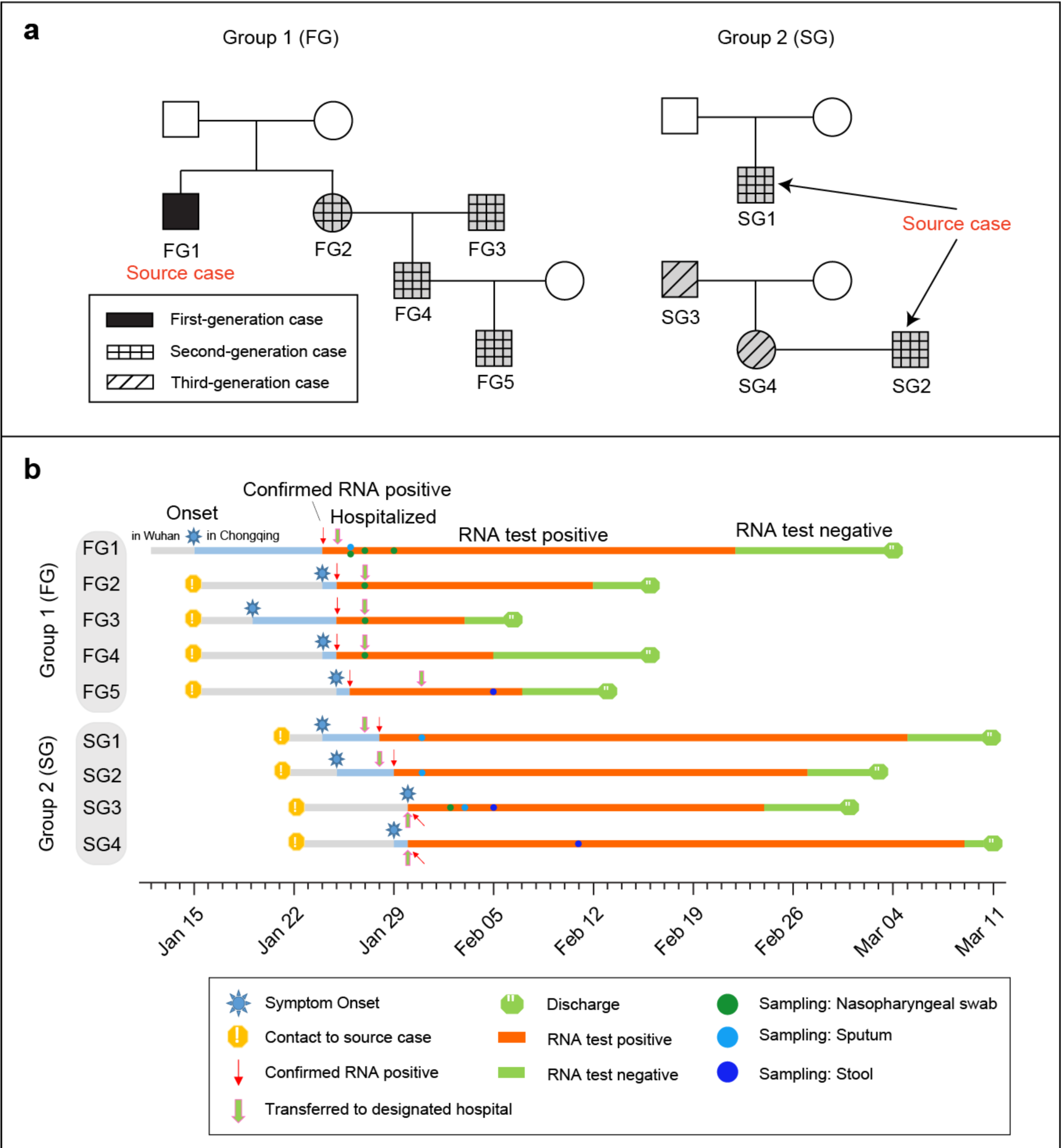
Details of participants and its related samples collected in this study. (**a**) The family trees of the patients with COVID-19. A total 9 cases could assign to two different single-source infection groups. Each groups contained one main family cluster. (**b**) The timeline of exposure history to index patients and the medical course of the participants.

**Figure 2.**
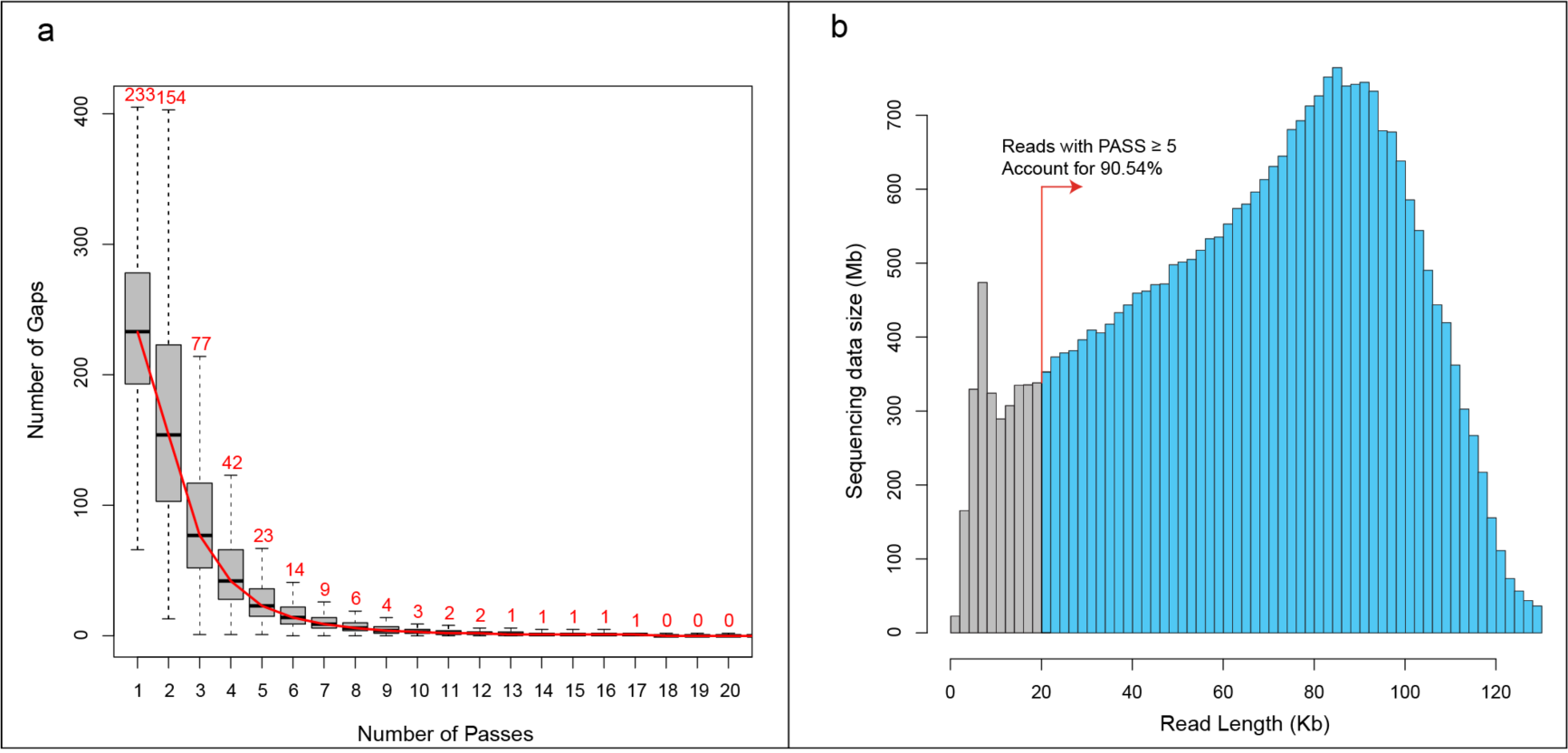
Quality assessments of circular consensus sequences (CCS). (**a)** Number of gaps in CCS under different circular sequencing passes. When pass was equal to 5, the number of gaps in CCS decreased to a stable value, which was close to completing self-correction. Meanwhile, accumulated length of subreads with CCS pass ≥ 5 accounted for 90.54% of total sequencing data (**b**).

### Evaluation of genetic diversity and evolutionary rate of quasispecies haplotypes

A total number of 282,866 qualified CCS (99.72%) contained a full length of spike gene, and could be merged into 25,490 quasispecies haplotypes (1,581-3,713; Table 1). Compared to an early ancestral strain (Wuhan-Hu-1, Jan 05, 2020; GenBank: MN908947.3), all haplotypes accumulated 8,978 single nucleotide variations (SNVs) and 6,458 amino acid changes. The average genetic distance of quasispecies was ~8.36 × 10^−4^, while the value of ST samples (9.14 × 10^−4^) was greater than that of NS (8.11 × 10^−4^) and SP samples (8.13 × 10^−4^) (Fig. 3a). We further observed that the percentage of missense mutations and nonsense mutations in ST samples was also higher than the other two sampling types, where the percentage of missense mutations of ST, NS, and SP was 78.76%, 75.90%, and 75.24%, the percentage of nonsense mutations was 4.59%, 3.53%, and 4.16%, respectively (**Supplementary Fig. 2**). Based on these mutations, the overall substitution rate of SARS-CoV-2 haplotypes was estimated to be 2.68-3.86 × 10^−3^ per site per year (Fig. 3b).

**Table 1.**
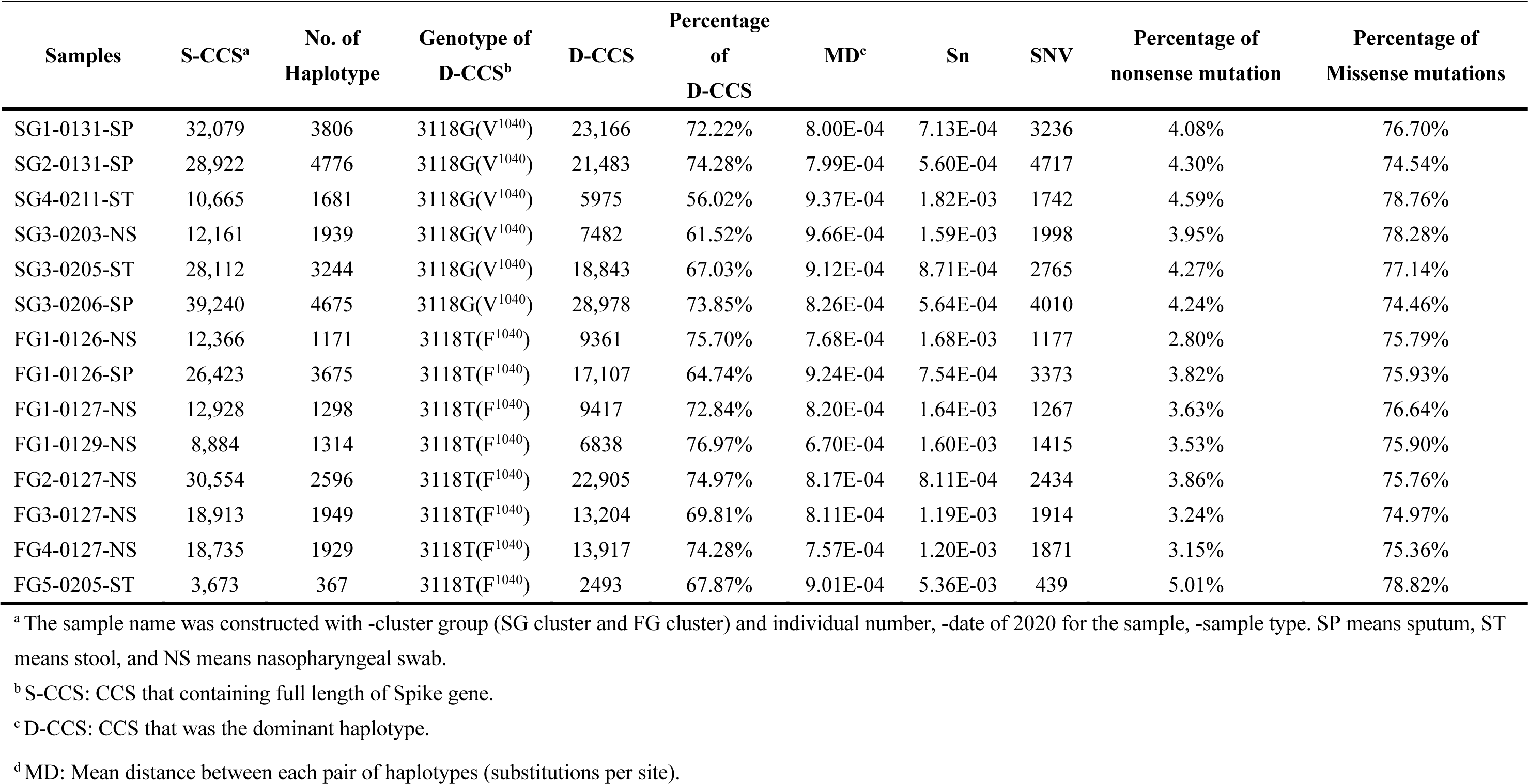
Information of SARS-CoV-2 quasispecies.

**Figure 3.**
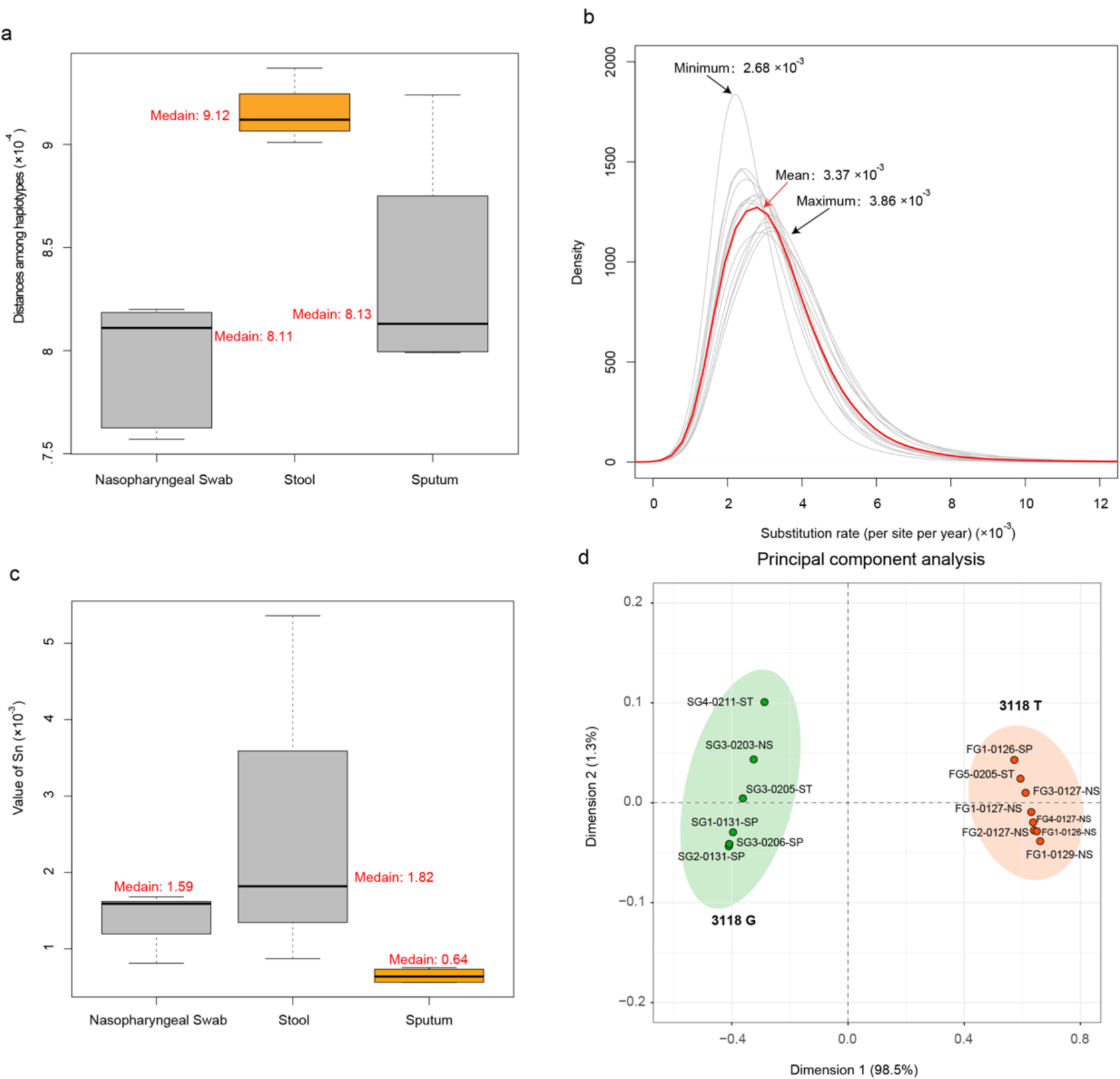
Analysis of the single base variations (SNVs) of quasispecies haplotypes. (**a**) Genetic distances of quasispecies haplotypes of SARS-Cov-2. Samples of stool showed obviously higher than other two sampling types. (**b**) Estimating the evolutionary rate at quasispecies level. The minimum, mean, and maximum values of 14 samples was 2.68×10^−3^, 3.37×10^−3^, and 3.86×10^−3^ substitution per site per year, respectively. (**c**) The standardized shannon entropy (Sn) of different sampling types. Sample from sputum had the lowest value. (**d**) Principal component analysis based on haplotype variations. The SNV G3118T as the first dimension could fully distinguish the two single-source infection groups.

### Identification of a novel quasispecies structure with the form of ‘unique dominant + numerous minors’

The quasispecies structure of all samples revealed the same style that unique dominant haplotypes with extremely high abundance of 56.02-76.97% (mean: 70.15%) combined with a huge number of minor haplotypes with extremely low abundance of < 5% (mean: < 0.1%) (**Supplementary Fig. 3**). The special quasispecies structure made a low quasispecies complexity that the overall Sn value was 1.35 × 10^−3^. For details, SP revealed significantly lower than other two sampling types (Chi-square test: p-value = 0.06), where the mean Sn value is 0.64 × 10^−3^, 1.59 × 10^−3^, and 1.82 × 10^−3^ for SP, NS, and ST, respectively (Fig. 3c). Two single-source groups contained different dominant haplotypes, where FG harbored an SNV 3118G>T and SG showed the same with the reference strain with 3118G (Table 1). The principal component analysis showed that the dimension of 3118G>T could be fully classified the samples into two groups and explained more than 98.5% of variances (Fig. 3d and **Supplementary Fig. 4**).

**Figure 4.**
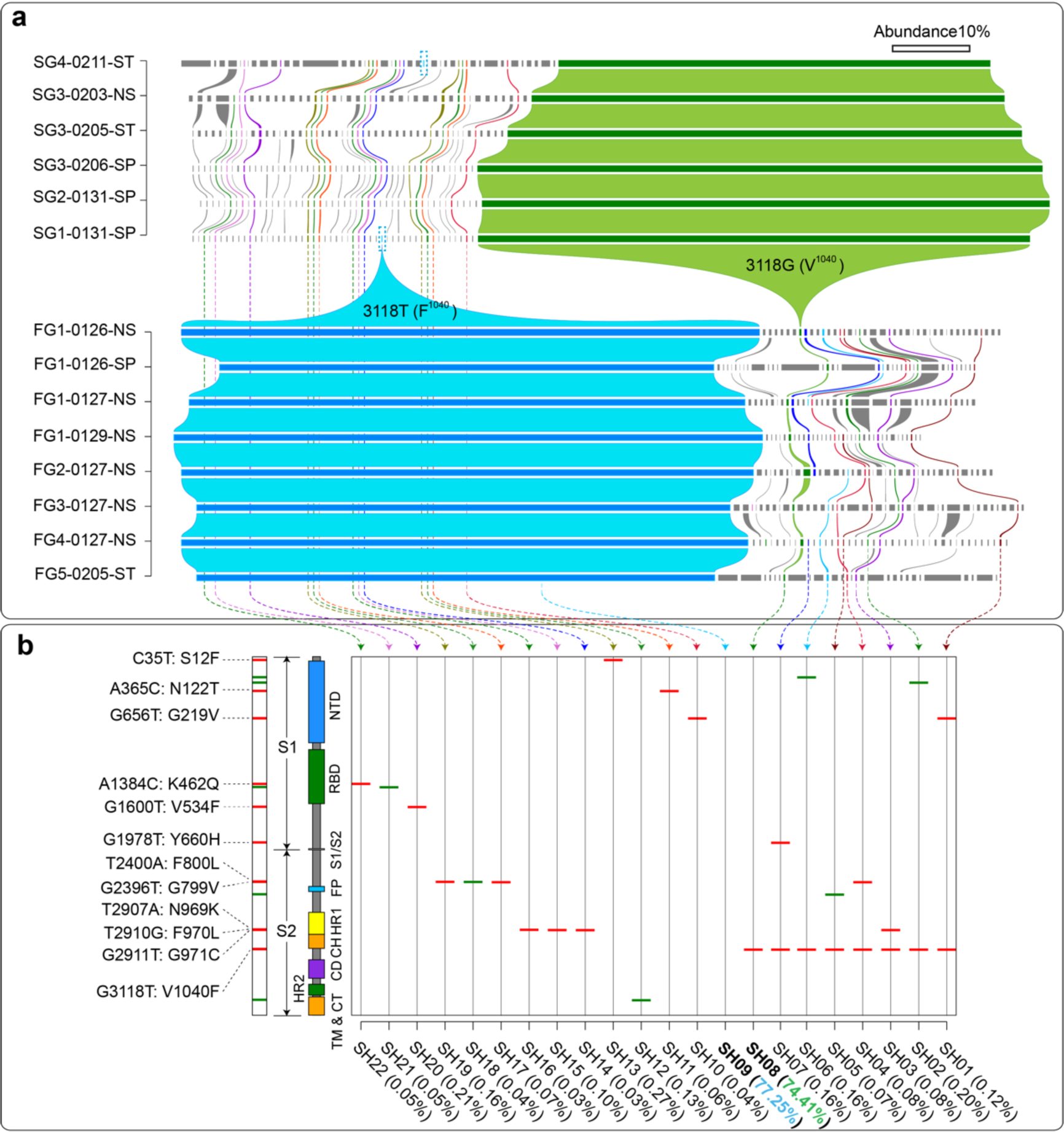
Content distribution and dynamics of the top 30 abundant quasispecies haplotypes. (**a**) Abundance distribution of quasispecies haplotypes in each sample. Each row represented one sample. Each rectangle represented a haplotype, and the width reflected its relative abundance. Only the top 30 abundant quasispecies haplotypes in any samples were considered. Colored haplotypes existed in at least six samples, and was defined as shared haplotypes (SH). Only ~6% minor haplotypes could be found as SH, indicating its high transmission bottleneck. Between any two adjacent samples, the same haplotype was connected by a polygon. All samples revealed the same quasispecies structure that one dominant haplotype followed by numerous minor haplotypes with low abundance. Two single infectious source groups contained two different dominant haplotypes. (**b**) Mutations on 22 shared haplotypes. Each line was an SH, and the red lines were the missense mutations and the green lines were synonymous mutations. All cumulative variations and its amino acid changes were showed in the left. Each functional domain was marked by rectangle. NTD, N-terminal domain; RBD, receptor-binding domain; S1/S2, protease cleavage site; FP, fusion peptide; HR1, heptad repeat 1; CH, central helix; CD, connector domain; HR2, heptad repeat 2; TM, transmembrane domain; CT, cytoplasmic tail. The name of SH were listed at the bottom, and the maximum abundant values were also listed in brackets.

### Observing the dynamics of quasispecies: high transmission bottleneck of minor haplotypes, and potentially multi-host origin of new dominant haplotypes

For the top 30 most abundant quasispecies of each sample, the dominant haplotypes 3118G and 3118T, together with other 20 minor ones could be shared by at least six samples. The shared haplotypes (SH) could be stably transmitted and survived between hosts or at different infection stage within hosts (Fig. 4a). A huge transmission bottleneck was observed for the minor haplotypes, where only ~ 5.66% (20/353) could be treated as SH. Dominant haplotype 3118G was the same with the reference early Wuhan strain (ancestor), and maintained the most stable existence (shared by all 14 samples). However, the abundance of 3118G decreased rapidly in FG familial cluster and became the minor haplotypes with a low abundance of 0.11 to 0.80% during spread. Instead, haplotype 3118T had replaced the ancestral 3118G as the major one in the first-generation case FG1 and further delivered to other cases of FG, while it was just a dust like minor haplotype in two SG samples (SG4-0211-ST: 0.009% and SG2-0131-SP: 0.003%). Compared with the published genome sequences (by August, 2020), the 3118G>T was a novel SNV resulting in Valine to Phenylalanine amino acid changes (V1040F) in S2 domain (Fig. 4b and **Supplementary Fig. 5**). By simulating the three-dimensional model of spike protein, we observed the mutation of V1040F may further change its corresponding R-group (**Supplementary Figs. 6 and 7**).

**Figure 5.**
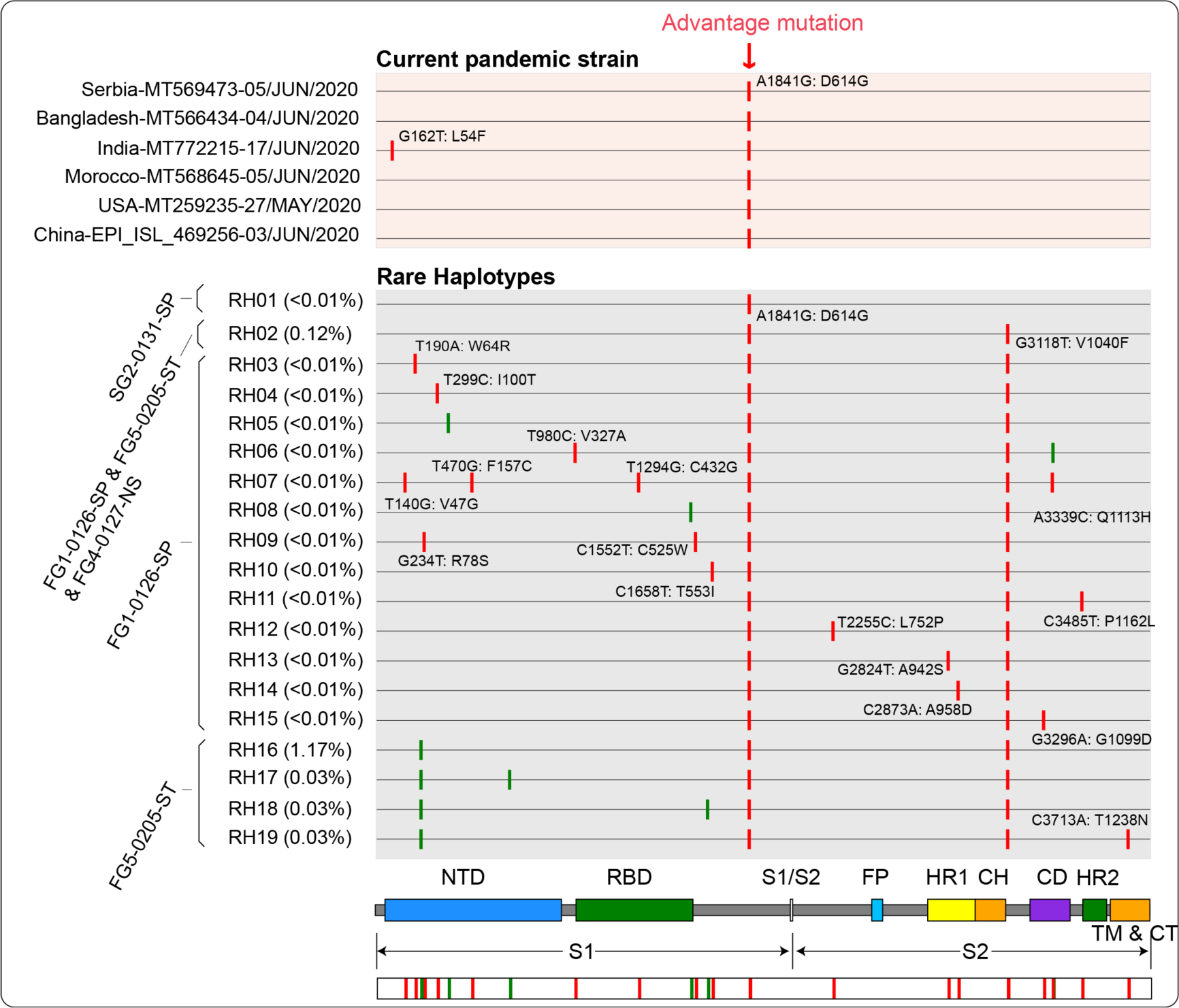
Haplotypes with the current pandemic mutations of 1841A>G (D614G). Haplotypes with red background were current pandemic strains collected from six countries, all which contained the fitness mutation of 1841A>G (D614G). The 1841G (G^614^) genotype had already existed in some SH of this study. Please refer to the legends of Fig. 4 for the meaning of each element in this Figure. FG cases (with 3118T dominant haplotype) had obtained the 1841G (G^614^) likely through viral mutations rather than infection (all with 1841G-3118T).

**Figure 6.**
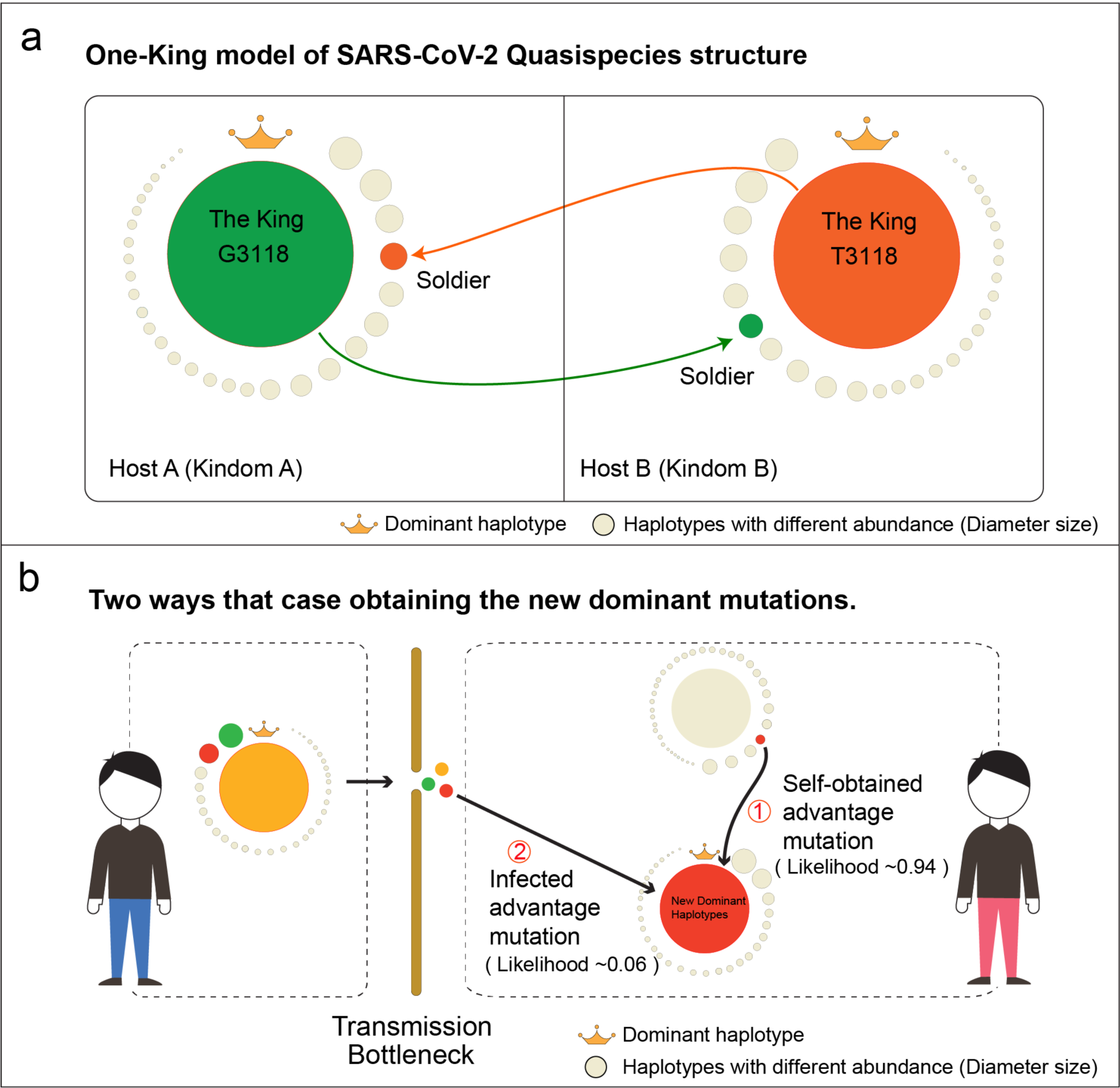
Models of the quasispecies structure and origin of dominant haplotypes. (**a**) The ‘One-King’ model of SARS-CoV-2 Quasispecies structure. All quasispecies haplotypes in a host were just like all people in an ancient kingdom. There can only be one king in a kingdom, with extremely great power (Unique dominant haplotype with extremely high abundance). In the kingdom, any capable soldier or normal person had a chance to become the king under some situations (Minor haplotypes with advantage mutations could potentially become the dominant one). (**b**) **Two ways that cases obtaining new dominant haplotype**. Due to the special quasispecies structure, the minor haplotypes had a high transmission bottleneck, and only ~6% could show stable transmission. For the cases with a new epidemic virus strain, the roughly likelihood of obtaining it through infection was estimated as 0.06, while the likelihood of self-obtained by mutating within host was 0.94. Our results indicated that new epidemic haplotypes were likely multi-host origin and could be emerged in different cases by self-obtaining.

Except for SH, most haplotypes (99.38%) were rare haplotypes (RH) that shared by less than five samples and with extremely low abundance, where 80.16% were singletons. These RH seemed to have no competitive advantage and survive as minor quasispecies in the early samples (mean abundance < 0.1%). However, we cannot ignore the existence of the SH, because it may become a dominant haplotype under certain conditions and cause a pandemic. The RH with mutation of 1841A>G (D614G) was currently prevalent in newly infected patients worldwide (Fig. 5). The mutation of 1841A>G revealed a high-frequency and was shared by 19 haplotypes, and most of them (18/19) were emerged by self-mutating of dominant haplotype 3118T (Fig. 5). The epidemic haplotype 1841G (G^614^) has already existed in early cases of our study, indicating the fitness haplotype may be multi-host origin. Due to high transmission bottleneck, cases obtained new advantage mutations were likely by self-mutating rather than transmission.

Based on our findings, we proposed a ‘One-King’ model to reflect the special quasispecies structure of SARS-CoV-2 (Figure 6a), and also inferred the high possibility of multi-host origin of the advantage mutations or haplotypes (Figure 6b). Finally, we extracted mainly two kinds of potentially fitness mutations within the SH and RH. One was high-abundant mutations on the top 30 abundant haplotuypes and the other was high frequency mutations shared by at least 15 haplotypes, which was separately found 225 and 337 missense mutations in different functional domains (**Supplementary Tables 3 and 4**).

## Discussion

Two infection chains with nine first-to third-generation cases made it possible to study the quasispecies and its dynamics during transmission (Fig. 1). SARS-Cov-2 revealed a special quasispecies composition that one dominant haplotype combined with numerous minor haplotypes. We not only discovered a novel advantage haplotype, but also realized that minor quasispecies were also worthy of attention, especially the minor haplotypes with high-abundant or high-frequent genetic variations. These minor ones may potentially exhibit a strong fitness and grow into the dominant one. However, due to the high transmission bottleneck, source hosts or cases obtained such advantage haplotypes or mutations were likely self-obtained from virus mutations rather than obtained by transmission. Our findings suggested its potential multi-host origin of an epidemic strain. Moreover, we modeled the special quasispecies structure as ‘One-King’, and it could explain incomprehensible phenomenon at the strain genome level, such as limited cumulative mutations ^14–17^ and low evolutionary rate ^25–27^. The current study could deepen our understanding of the underlying genetic mechanism on the fitness changes of SARS-Cov-2.

High-quality haplotype sequences lay the foundation for the subsequent analysis and its interpretation. More than 98% virus genomes contained the identical sequences to the amplified primers, indicating the CCS results have no primer bias (**Supplementary Fig. 1**). We obtained an average 27,000 quasispecies sequences for each sample, and the number was more than the SARS-Cov quasispecies study using traditional sanger methods (28 spike sequences from 19 patients) ^28^ and results from the recent HCV quasispecies study using the same SMRT platform (average 8,130 CCS for each samples with pass > 5) ^29^. More than 80,000 cumulative SNVs facilitated us to calculate the evolutionary rate at quasispecies level. The average substitution rate of SARS-Cov-2 was estimated as 2.68-3.86×10^−3^ per site per year (Fig. 3b), which was larger than its recent estimation value of 0.99-1.83×10^−3^ at the consensus genome level ^25–27^. Compared with other *betacoronaviruses*, the evolutionary rate of SARS-Cov-2 was similar to that of SARS-Cov (1.16 – 3.30 ×10^−3^ and 1.67 – 4.67×10^−3^ for non-synonymous and synonymous mutations, respectively)^30^, but slightly larger than MERS-Cov (1.23-2.04×10^−3^) ^31^.

We observed a special quasispecies structure of SARS-Cov-2 that unique dominant haplotypes with abundance of 56.02-76.97% combined with numerous minor haplotypes with mean abundance of < 0.1%, which seemed like ‘master + dust’ (**Supplementary Fig. 3**). The proportion of main quasispecies was much higher than that of SARS-Cov ^28^ and other RNA virus like HBV and HCV ^32,33^. Two single-source infection groups had different dominant haplotypes, which was consistent with two independent infection chains. The dominant haplotype of SG (V^1040^) was the same with the strain from early stage of the epidemic. However, a novel haplotype with F^1040^ replaced the V^1040^ became the dominant one in FG, while the V^1040^ haplotype still existed as minor form (Figs. 4a and b). All indicated that the dominant haplotype and minor haplotype could be switched under some situations. The same phenomenon was also found in rare haplotypes (RH). The currently epidemic strain 1841G (G^614^) ^21^ ^22^ has already existed in our early samples as the minor type (Fig. 5). Although the novel dominant genotype F^1040^ may have changed the structure of spike protein to obtain a fitness advantage (**Supplementary Figs. 5-7**), it has not spread as widely as G^614^, which may be due to timely clinical testing and isolation treatment.

According to these findings, we termed and used the ‘One-King’ model to represent this special quasispecies structure. All quasispecies haplotypes in a host were just like all people in an ancient kingdom. There can only be one king in a kingdom, with extremely great power (Unique dominant haplotype with extremely high abundance). In the kingdom, any capable soldier or normal person had a chance to become the king under some situations (Minor haplotypes with advantage mutations could potentially become the dominant one) (Fig. 6a). The ‘One-King’ model could explain some incomprehensible phenomena at strain genome level. The reason why SARS seemed to have only limited cumulative mutations and a small evolutionary rate was because limited dominant haplotypes determined the consensus genome sequences, while ignoring a large amount of minor haplotypes and its mutations.

Our results observed the possibility that some advantage minor haplotypes becoming the dominant one. However, such minor haplotypes were likely self-obtained from virus rather than from transmission, because the ‘One-King’ structure was more conducive to the spread of major haplotypes, but likely caused a transmission bottleneck for minor ones. That was why only a small proportion of minor haplotypes (< 6%) could be shared by cases in the same infection chain (Fig. 4), and also all hosts with dominant haplotype 3118T (F^1040^) obtained the advantage mutation A1841G (G^614^) by self-mutated (Fig. 5). The high likelihood of self-obtained (~0.94) suggested that the dominant haplotype may be of multi-host origin (Fig. 6b), which was the possible reason why the dominant mutation of D614G appears in different cases with geographical isolation worldwide (Fig. 5). Although this kind of fitness mutations emerged in minor haplotypes, their frequency was relative high, indicating a selective advantage.

Due to the special structure and short evolution time, the genetic distance and complexity of SARS-CoV-2 quasispecies revealed greatly low. Nevertheless, quasispecies features could still be showed different between sampling types. Samples from stools had a larger genetic distance and more nonsense mutations and missense mutations than nasopharyngeal swab, and sputum (Fig. 3a and **Supplementary Fig. 2**), indicating stool samples contained relatively more inactive virus. In other aspects, the standard entropy (Sn) of sputum samples was the lowest (Fig. 3c), indicating its haplotype composition was more dispersed than other two types, and its minor haplotypes may have a larger transmission bottleneck.

Our study had some limitations. Firstly, due to technical limitations, it was currently difficult to obtain a haplotype with full-length of genome (~29Kb), and the spike gene (~4Kb) may be partially reflected the genome-wide quasispecies information. Secondly, insufficient number of samples made it difficult to observe the dynamics in a relatively large scope such as the intermediate transition state. Thirdly, the current study only focuses on SNVs, and other mutation types such as insertions, deletions, and structural variations need to be further studied.

The ‘One-King’ structure of SARS-CoV-2 quasispecies reflected a novel quasispecies genetic fitness mechanism. Behind the dominant haplotype, there were countless minor fitness haplotypes that could be survived as dominant one in different hosts. Although these haplotypes may be suppressed and unable overcome the transmission bottleneck, the advantage mutations within them could be high-frequently obtained by virus mutations. Future molecular epidemiologic investigations, clinical interventions and vaccine designs need to consider the potential advantage SARS-CoV-2 quasispecies and its related mutations, which are missed by routine single concensus genotype ^34^ (**Supplementary Tables 3 and 4**).

## Methods

### Participants in the study

Nine patients with laboratory-confirmed COVID-19 were included in this study. All of them were admitted to Chongqing Public Health Medical Center (CPHMC), the designated hospital for COVID-19 treatment in Chongqing centre area. According to the epidemic history, all patients were divided into two single-source infection cluster groups, SG cluster and FG cluster group (Fig. 1a). The first group contained a familial cluster containing five SARS-CoV-2 infected patients, including one first-generation case infected in Wuhan (patient: FG1), and four second-generation cases (FG2, FG3, FG4, and FG5) infected by FG1 in Chongqing, China (Fig. 1b). The first case FG1 (a 70-year-old male) was a local resident in Wuhan, and came to Chongqing to visit his younger sister’s family (FG2, FG2’s husband: FG3, FG2’s son: FG4, FG2’s grandson: FG5) in Jan 15 of 2020 by taking a train and then they lived together from that day. It was on that day he presented a symptom of fever (highest temperature 37.7 °C), cough, and took antipyretic himself. He went to clinic on Jan 21 since no relief of symptoms and was test positive of SARS-CoV-2 in Jan 24 and then he was transferred to the designated hospital CPHMC. His younger sister FG2 (60-year-old), the husband FG3 (62-year-old, P3), the son FG4 (36-year-old) and the grandson FG2 (10-year-old) start to present fever and or cough symptom with highest temperature 39.3 °C, 38 °C 37.5 °C, 38 °C since Jan 24, Jan19, Jan24, and Jan25, respectively. All FG2, FG3, FGP4 and FG5 were quarantined as close contacts since FG1 was conformed as second-generation case, and received test in Jan 25 and Jan 26, the results came positive and then they were transferred to CPHMC also in Jan 27 and Jan 31. CT scan showed classical diffuse ground-glass opacity in both sides of lung for all five patients, and both they received Lopinavir/Ritonavir plus interferon-α antiviral treatment. Additionally, FG1, FG2, FG3 was given oxygen support though nasal cannula. FG1 had developed severe illness with the condition of 15-year hypertension. All of them recovered and discharged from hospital after a period therapy.

The second group contains four second- or third-generation cases infected by the same first-generation case, where one patient (SG1: 2^nd^) from an unknown family and three patients from the same family (SG2: 2^nd^; SG3: 3^rd^; SG4: 3^rd^) (Fig. 1b). The first generation case returned to Chongqing from Wuhan on January 20 and next had a dinner with two friends (SG1 and SG2) on January 21. SG1 and SG2 presented a symptom of fever, cough, and took antipyretic by self, and received positive test result in Jan 23, Jan 24 and Jan 25. Through direct contact, two family members of SG2 were also infected and revealed viral RNA positive on 19 Jan, including SG2’s wife SG4 and SG4’s mother SG3. According to the order of contact, SG1 and SG2 were considered as second cases, while SG3 and SG4 were third cases. Fourteen samples, include nasopharyngeal swab (NS), sputum (SP), and stool (ST), were collected from these 9 patients. The time points of samples were descripted in Fig 1b. All of them recovered and discharged from hospital after a period therapy. All the samples were from our previous study ^35^, which was approved by the ethics committee of CPHMC (document no. 2020-002-01-KY, 2020-003-01-KY) and conducted in accordance with Declaration of Helsinki principles. Written informed consent was obtained from each subject.

### Extraction and amplification of the spike gene

Viral RNA was extracted by using the EZ1 virus mini kit of QIAGEN Company in Germany, and then 8μL of each was reverse transcribed into cDNA by using the commercial company's reverse transcription kit (TAKARA No. 6110A). The full-length spike gene (3882 nt) was further amplified by a pair of RT-PCR primers designed by online methods from NCBI. The forward primer was: 5 '-barcod-AGGGGTACTGCTGTTATGTCT-3'), and the reverse primer was: 5'-barcod-GCGCGAACAAAATCTGAAGG-3'. We used 50 μl PCR system to conduct the PCR reaction, and the system includes 5 μL SARS-CoV-2 cDNA, 2.5μl forward and reverse primer (10μmol/L), 25μl Q5^®^ High-Fidelity 2X Master Mix (NEB No. M0492S), complement volume with Nuclease-Free Water (Thermofisher). PCR reaction conditions: denaturation 98°C/30s, circulation (35 times) 98°C/10s, 54°C/30s, 72°C/ 3min; extension 72°C/6min. The PCR product was electrophoretic in 0.8% agarose gel to verify whether the product was successfully amplified. Next, using Ampure PB beads of Pacific Biosciences company in the USA for pcr products size selection: First, the Ampure PB beads were diluted to a 35% concentration using elution buffer; Then, the diluted Ampure PB beads were used for enrichment fragments larger than 3Kb (Target fragment was ~4 Kb), the volume of the diluted Ampure PB beads is about 3 times of the sample volume. After size selection, full-length DNA of S gene was quantified using The Qubit 2.0 Fluorometer and Qubit dsDNA HS Assay Kit (Thermofisher No. Q32851).

Due to sample quality or load of viral RNA, not all three sampling types could be successfully amplified the target segment. Seven samples could be obtained the next-generation sequencing reads from metagenomic sequencing, which was used to access the amplification bias by BWA software ^36^ (**Supplementary Fig. 1**). Finally, a total number of 14 samples passed the quality control and screened for further analysis (**Supplementary table 1**). The sample name reflected three aspects of information including patients, sampling date, sources of nucleic acid. For example, FG1-0126-NS meant viral RNA was extracted form nasopharyngeal swabs of FG1 patients on January 26, 2020.

### Single-molecule real-time (SMRT) sequencing of the target segments

About 100ng full-length DNA of spike gene was used for library construction. For briefly, the full-length cDNA was subjected to damage repair, end repair/A-tailing, and ligation of the SMRT adapter and unique label for each sample. The primers and DNA-binding polymerase were combined to generate a complete SMRT bell library. After qualitatively analyzing the library, the PacBio Sequel I platform was used for sequencing according to the effective concentration of the library and the data output requirements. We applied SMRT Link software to obtain the circular consensus sequences (CCS), and only CCS with over 5 full-passes were considered for further analysis (Fig. 2).

### Data processing

After performing the quality control and removing the low-quality subreads (minimum predicted accuracy of 90%), we totally obtained 26.97 Gb of subreads for all 14 samples (**Supplementary table 1**). Our results showed that the number of CCS gaps decreased rapidly and tended to be stable when pass equaled 5 (Fig. 2a). Further statistics showed that such reads account for ~90% of total sequencing data (Fig. 2b and **Supplementary Table 2**). The spike gene (S gene) of strain Wuhan-Hu-1 (Extracted from patients with early infection in Wuhan; GenBank accession number MN908947.3, it was submitted in January 5, 2020)^37^ was used as reference to detect single nucleotide variations (SNVs). We applied BLAST^38^ software to perform alignment between CCS and the reference, and then obtained all the CCS results containing full length of spike gene. A total number of 282,866 CCS were finally identified (Table 1). The complexity and diversity of quasispecies were detected by calculating standardized Shannon entropy (Sn) and mean genetic distance (d), respectively (Fig. 3 and **Supplementary Fig. 2**). The former formula is Sn = – Σi (pi*Inpi) / InN, where pi was the frequency of the top abundance quasispecies, and N was the total number of quasispecies haplotypes. Meanwhile, SNV information were extracted using PERL scripts based on the BLAST results. CCS with the same haplotype was further clustered into the same quasispecies haplotype, and its abundance was further calculated by PERL scripts (**Supplementary Fig. 3**). The R program was applied to perform Principle component analysis (Fig. 3 and **Supplementary Fig. 2**). The three-dimensional ribbon models for the spike protient was prepared by UCSF Chimera^39^ based on the entry 6VSB in Protein Data Bank ^40^ (**Supplementary Figs. 5 and 6**). MEGAX^41^ was used to calculate the mean genetic distance and carry out the phylogenetic analysis. The potential fitness mutations were annotated and extracted by using in-house PERL scripts (**Supplementary tables 3 and 4)**.

## Data availability

All other data are included in the supplemental information or available from the authors upon reasonable requests. Source data are provided with this paper.

## Acknowledgements

The current study was supported by Chongqing Health Commission COVID-19 Project (2020NCPZX01), TMMU COVID-19 Project (2020XGBD09), Youth Talent Medical Technology Program of PLA (17QNP010), Chinese Key Project Specialized for Infectious Diseases (2018ZX10723203), and the TMMU key project for medical research (2018XYY10). We thank for the supports the Youth Talent Program from Third Military Medical University (W. Tan and F. Sun) and the Academy of Medical Sciences Newton International Fellowship (Tan W). We also thank all patients that participated in this study.

## Author contributions

Drs Deng, W. Tan and Chen had full access to all of the data in the study and take responsibility for the integrity of the data and the accuracy of the data analysis. Concept and design: Deng, W. Tan, Chen; Acquisition, analysis, or interpretation of data: Sun, Wang, Dan, S. Tan, Xu; Sample preparation: J. Zhang, Lu, Z. Tan, Xiang, Zhou, He, Wan, W. Zhang; Drafting of the manuscript: Deng, Sun, W. Tan; Critical revision of the manuscript for important intellectual content: W. Tan; Supervision: Deng.

## Competing interests

The authors declare no competing interests.

## Notes

### Competing Interest Statement

The authors have declared no competing interest.

## References

1. Snijder, E.J. et al. Ultrastructure and origin of membrane vesicles associated with the severe acute respiratory syndrome coronavirus replication complex. J Virol 80, 5927–40 (2006).

2. Han, H.J. et al. Bats as reservoirs of severe emerging infectious diseases. Virus Research 205, 1–6 (2015).

3. Molecular evolution of the SARS coronavirus during the course of the SARS epidemic in China. Science 303, 1666–9 (2004).

4. Zhong, N.S. et al. Epidemiology and cause of severe acute respiratory syndrome (SARS) in Guangdong, People's Republic of China, in February, 2003. Lancet 362, 1353–8 (2003).

5. Ge, X.Y. et al. Isolation and characterization of a bat SARS-like coronavirus that uses the ACE2 receptor. Nature 503, 535–+ (2013).

6. Zumla, A., Hui, D.S. & Perlman, S. Middle East respiratory syndrome. Lancet 386, 995–1007 (2015).

7. Zaki, A.M., van Boheemen, S., Bestebroer, T.M., Osterhaus, A.D.M.E. & Fouchier, R.A.M. Isolation of a Novel Coronavirus from a Man with Pneumonia in Saudi Arabia. New England Journal of Medicine 367, 1814–1820 (2012).

8. Oh, M.D. et al. Middle East respiratory syndrome: what we learned from the 2015 outbreak in the Republic of Korea. Korean Journal of Internal Medicine 33, 233–246 (2018).

9. Zhu, N. et al. A Novel Coronavirus from Patients with Pneumonia in China, 2019. New England Journal of Medicine 382, 727–733 (2020).

10. Kasibhatla, S.M., Kinikar, M., Limaye, S., Kale, M.M. & Kulkarni-Kale, U. Understanding evolution of SARS-CoV-2: A perspective from analysis of genetic diversity of RdRp gene. J Med Virol (2020).

11. Zhou, P. et al. Discovery of a novel coronavirus associated with the recent pneumonia outbreak in humans and its potential bat origin. bioRxiv, 2020.01.22.914952 (2020).

12. Mulangu, S. et al. A Randomized, Controlled Trial of Ebola Virus Disease Therapeutics. New England Journal of Medicine 381, 2293–2303 (2019).

13. Wang, M. et al. Remdesivir and chloroquine effectively inhibit the recently emerged novel coronavirus (2019-nCoV) in vitro. Cell Res 30, 269–271 (2020).

14. Phan, T. Genetic diversity and evolution of SARS-CoV-2. Infect Genet Evol 81, 104260 (2020).

15. Pachetti, M. et al. Emerging SARS-CoV-2 mutation hot spots include a novel RNA-dependent-RNA polymerase variant. J Transl Med 18, 179 (2020).

16. Benvenuto, D. et al. Evolutionary analysis of SARS-CoV-2: how mutation of Non-Structural Protein 6 (NSP6) could affect viral autophagy. Journal of Infection 81, E24–E27 (2020).

17. Xiaolu, T. et al. On the origin and continuing evolution of SARS-CoV-2. National Science Review (2020).

18. Vignuzzi, M., Stone, J.K., Arnold, J.J., Cameron, C.E. & Andino, R. Quasispecies diversity determines pathogenesis through cooperative interactions in a viral population. Nature 439, 344–8 (2006).

19. Zhou, P. et al. A pneumonia outbreak associated with a new coronavirus of probable bat origin. Nature 579, 270–273 (2020).

20. Becerra-Flores, M. & Cardozo, T. SARS-CoV-2 viral spike G614 mutation exhibits higher case fatality rate. Int J Clin Pract, e13525 (2020).

21. Kim, S.J., Nguyen, V.G., Park, Y.H., Park, B.K. & Chung, H.C. A Novel Synonymous Mutation of SARS-CoV-2: Is This Possible to Affect Their Antigenicity and Immunogenicity? Vaccines (Basel) 8(2020).

22. Korber, B. et al. Tracking Changes in SARS-CoV-2 Spike: Evidence that D614G Increases Infectivity of the COVID-19 Virus. Cell (2020).

23. Li, W. et al. Angiotensin-converting enzyme 2 is a functional receptor for the SARS coronavirus. Nature 426, 450–4 (2003).

24. Raj, V.S. et al. Dipeptidyl peptidase 4 is a functional receptor for the emerging human coronavirus-EMC. Nature 495, 251–4 (2013).

25. Li, X. et al. Transmission dynamics and evolutionary history of 2019-nCoV. J Med Virol 92, 501–511 (2020).

26. Nie, Q. et al. Phylogenetic and phylodynamic analyses of SARS-CoV-2. Virus Research, 198098 (2020).

27. Li, X. et al. Evolutionary history, potential intermediate animal host, and cross-species analyses of SARS-CoV-2. J Med Virol 92, 602–611 (2020).

28. Xu, D., Zhang, Z. & Wang, F.S. SARS-associated coronavirus quasispecies in individual patients. N Engl J Med 350, 1366–7 (2004).

29. Yamashita, T. et al. Single-molecular real-time deep sequencing reveals the dynamics of multi-drug resistant haplotypes and structural variations in the hepatitis C virus genome. Sci Rep 10, 2651 (2020).

30. Zhao, Z. et al. Moderate mutation rate in the SARS coronavirus genome and its implications. BMC Evol Biol 4, 21 (2004).

31. Tong, Y.G. et al. Genetic diversity and evolutionary dynamics of Ebola virus in Sierra Leone. Nature 524, 93–6 (2015).

32. Tsukiyama-Kohara, K. & Kohara, M. Hepatitis C Virus: Viral Quasispecies and Genotypes. Int J Mol Sci 19(2017).

33. Zhang, A.Y. et al. Deep sequencing analysis of quasispecies in the HBV pre-S region and its association with hepatocellular carcinoma. J Gastroenterol 52, 1064–1074 (2017).

34. Andersen KG, Rambaut A, Lipkin WI, Holmes EC, Garry RF. The proximal origin of SARS-CoV-2. Nat Med 2020. https://doi.org/10.1038/s41591-020-0820-9

35. Tan, W. et al. Viral Kinetics and Antibody Responses in Patients with COVID-19. medRxiv, 2020.03.24.20042382 (2020).

36. Li, H. & Durbin, R. Fast and accurate short read alignment with Burrows-Wheeler transform. Bioinformatics 25, 1754–60 (2009).

37. Wu, F. et al. A new coronavirus associated with human respiratory disease in China. Nature 579, 265–269 (2020).

38. Chen, Y., Ye, W., Zhang, Y. & Xu, Y. High speed BLASTN: an accelerated MegaBLAST search tool. Nucleic Acids Res 43, 7762–8 (2015).

39. Pettersen, E.F. et al. UCSF Chimera--a visualization system for exploratory research and analysis. J Comput Chem 25, 1605–12 (2004).

40. Wrapp, D. et al. Cryo-EM structure of the 2019-nCoV spike in the prefusion conformation. Science 367, 1260–1263 (2020).

41. Kumar, S., Stecher, G., Li, M., Knyaz, C. & Tamura, K. MEGA X: Molecular Evolutionary Genetics Analysis across Computing Platforms. Mol Biol Evol 35, 1547–1549 (2018).

