## Supplementary for "SARS-CoV-2 Quasispecies provides insight into its genetic dynamics during infection"

### Supplementary tables and figures

**Supplementary Table 1 Information of screened cases and its related samples.**

| Group ID | Family ID | Sample ID | Patient | Infectious generation | Sampling date | Sample type | CT[a] |
|---|---|---|---|---|---|---|---|
| FG | 1 | FG1-0126-NS | FG1 | 1st | January 26, 2020 | Nasopharyngeal Swabs | 24.1 |
| | | FG1-0127-NS | | | January 27, 2020 | Nasopharyngeal Swabs | 22.9 |
| | | FG1-0129-NS | | | January 29, 2020 | Nasopharyngeal Swabs | 19.7 |
| | | FG1-0126-SP | | | January 26, 2020 | Sputum | 16.3 |
| | | FG2-0127-NS | FG2 | 2nd | January 27, 2020 | Nasopharyngeal Swabs | 26.2 |
| | | FG3-0127-NS | FG3 | 2nd | January 27, 2020 | Nasopharyngeal Swabs | 28.8 |
| | | FG4-0127-NS | FG4 | 2nd | January 27, 2020 | Nasopharyngeal Swabs | 23.4 |
| | | FG5-0205-ST | FG5 | 2nd | February 05, 2020 | Stool | 20.4 |
| SG | 2 | SG1-0131-SP | SG1 | 2nd | January 31, 2020 | Sputum | 18.3 |
| | 3 | SG2-0131-SP | SG2 | 2nd | January 31, 2020 | Sputum | 11.9 |
| | | SG3-0203-NS | SG3 | 3rd | February 03, 2020 | Nasopharyngeal Swabs | 19.7 |
| | | SG3-0205-ST | | | February 05, 2020 | Stool | 19.7 |
| | | SG3-0206-SP | | | February 06, 2020 | Sputum | 13.7 |
| | | SG4-0211-ST | SG4 | 3rd | February 11, 2020 | Stool | 22.3 |

Note: [a] represented for cycle threshold for real-time PCR.

**Supplementary Table 2 Statistics of sequencing data**

| Samples | Sequencing Data Size(Gb) | Pass number of CCS | | | | | | | | | |
|---|---|---|---|---|---|---|---|---|---|---|---|
| | | ≥1 | ≥ 2 | ≥3 | ≥ 4 | ≥ 5 | ≥ 6 | ≥ 7 | ≥ 8 | ≥ 9 | ≥10 |
| SG1-0131-SP | 2.62 | 42,763 | 40,963 | 37,819 | 34,703 | 32,079 | 29,799 | 27,633 | 25,629 | 23,795 | 22,118 |
| SG2-0131-SP | 2.35 | 40,122 | 38,309 | 34,979 | 31,783 | 28,922 | 26,522 | 24,336 | 22,284 | 20,413 | 18,671 |
| SG4-0211-ST | 1.24 | 15,141 | 14,552 | 13,207 | 11,798 | 10,665 | 9,697 | 8,777 | 7,960 | 7,189 | 6,492 |
| SG3-0203-NS | 1.20 | 17,336 | 16,647 | 15,085 | 13,463 | 12,161 | 11,040 | 10,059 | 9,186 | 8,352 | 7,584 |
| SG3-0205-ST | 2.48 | 37,408 | 35,760 | 33,010 | 30,329 | 28,112 | 26,069 | 24,268 | 22,546 | 20,928 | 19,414 |
| SG3-0206-SP | 3.35 | 52,739 | 50,392 | 46,429 | 42,547 | 39,240 | 36,312 | 33,820 | 31,420 | 29,141 | 27,012 |
| FG1-0126-NS | 1.28 | 16,290 | 15,397 | 14,220 | 13,250 | 12,366 | 11,656 | 10,938 | 10,319 | 9,736 | 9,124 |
| FG1-0126-SP | 2.94 | 34,074 | 32,642 | 30,536 | 28,297 | 26,423 | 24,806 | 23,365 | 22,094 | 20,872 | 19,803 |
| FG2-0127-NS | 1.67 | 16,802 | 16,004 | 14,794 | 13,780 | 12,928 | 12,166 | 11,386 | 10,687 | 10,044 | 9,426 |
| FG1-0129-NS | 0.92 | 11,731 | 11,159 | 10,246 | 9,512 | 8,884 | 8,320 | 7,792 | 7,307 | 6,858 | 6,423 |
| FG2-0127-NS | 3.22 | 39,634 | 37,756 | 34,975 | 32,674 | 30,554 | 28,753 | 27,016 | 25,426 | 23,859 | 22,382 |
| FG3-0127-NS | 1.83 | 25,236 | 23,427 | 21,622 | 20,191 | 18,913 | 17,728 | 16,627 | 15,633 | 14,682 | 13,841 |
| FG4-0127-NS | 1.54 | 24,759 | 23,368 | 21,589 | 20,085 | 18,735 | 17,577 | 16,506 | 15,531 | 14,582 | 13,651 |
| FG5-0205-ST | 0.33 | 5,044 | 4,672 | 4,325 | 3,975 | 3,673 | 3,397 | 3,178 | 2,926 | 2,725 | 2,536 |

**Supplementary Table 3 Information of missense mutations with the top 30 abundant haplotypes of all samples.**

| Type | Domains | Nucleic acid variations | Amino acid variations |
|---|---|---|---|
| Shared Haplotypes (SH) | NTD | G656T; | G219V; |
| | RBD | A1384C; | K462Q; |
| | HR1 | T2910G; | F970L; |
| | Others | C35T;G1600T;G2396T;G3118T;T1978C;T2400A; | F800L;G799V;S12F;V1040F;V534F;Y660H; |
| Rare Haplotypes (RH) | NTD | A443C;A461G;A544G;A546T;A606T;A620G;A644G;A813T;A911G;C145T;C205T;C227T;C341T;C471A;C59T;C623T;C779T;C77T;C863T;C877T;C883T;C905T;G162T;G224A;G224C;G265T;G388T;G425A;G456T;G637T;G655A;G664T;G682T;G695C;G782A;G868C;G868T;G892C;G909T;T107C;T140G;T185G;T190A;T470C;T47C;T47G;T495G;T555G;T580G;T823C; | A222S;A260V;A288V;D215G;D228Y;D290H;D290Y;E154G;E298Q;F157L;F157S;F194V;F275L;G142D;G219S;G232A;G261D;G75A;G75D;G89C;H207R;H49Y;H69Y;K182E;K182N;K202N;K304R;L293F;L303F;L54F;N148T;N165K;N185K;P26L;P295S;Q271H;T114I;T208M;T20I;T302M;T76I;V130F;V16A;V16G;V213L;V36A;V47G;V62G;W152C;W64R; |
| | RBD | A1153C;A1162G;A1386C;C1055T;G1037A;G1082T;G1087A;G1099T;G1144T;G1211T;G1223T;G1297T;G1337T;G1397A;G1426T;T1057C;T1111G;T1129C;T1174C;T1327C;T1522C;T1544G;T1557G; | A352V;A363T;C361F;F377L;F392L;F515C;G404V;G446V;G476C;H519Q;K462N;N388D;R346K;R408I;R466K;S371A;S443P;T385P;V367F;V382L;V433F;W353R;Y508H; |
| | S1S2 | C2047T;G2048A;G2054T; | R683Q;R683W;R685L; |
| | HR1 | A2734G;A2858G;A2905G;C2812T;C2873A;C2947T;G2743T;G2824T;G2872T;G2912A;T2732C;T2813C; | A942S;A958D;A958S;G971D;L938F;L938P;N953S;N969D;R983C;T912A;V911A;V915F; |
| | CH | G2971A;G2995T;G3012T;T3002G; | G999C;L1001R;L1004F;V991M; |
| | CD | A3292G;A3400C;C3247T;C3299T;C3407T;G3296A;G3306T;G3320T;G3371T;T3265C;T3266G;T3284C;T3284G;T3307G; | F1089C;F1089L;F1095C;F1095S;F1103V;G1099D;G1124V;H1083Y;N1098D;N1134H;R1107M;T1100I;T1136I;W1102C; |
| | HR2 | A3514G;A3571G;A3575T;A3584C;C3521T;C3538A;G3554A | A1174V;E1195A;I1172V;I1179T;K1191E;K1191N;N1192I;Q1 |

| | | | |
|---|---|---|---|
| | | ;G3573T;T3536C; | 180K;R1185H; |
| | TM&CT | G3624T;G3650T;G3651T;G3672T;G3711C;G3716T;G3722T;G3735T;G3769T;T3640C;T3647C;T3701C;T3767G; | C1241F;D1257Y;F1256C;I1216T;K1245N;L1224F;L1234P;M1237I;Q1208H;S1239I;W1214R;W1217C;W1217L; |
| | Others | A1605T;A1841G;A2128G;A2200G;A2252G;A2328T;A2351G;A2432G;A2437G;A2605G;A2684T;A2702G;A3083T;A3122G;A3215G;A41T;C1640T;C1718T;C1796T;C1919T;C1993T;C2081T;C2102T;C2282T;C2293T;C2333A;C2333T;C2372T;C2434T;C2521T;C2636T;C2648T;C3026T;C3475A;C3484T;C964G;G1599T;G1600A;G1600T;G1749T;G1781T;G1843C;G1943T;G1966T;G1983C;G2000A;G2000C;G2006C;G2031T;G2101T;G2306A;G2314T;G2396A;G24C;G2700T;G3041C;G3043A;G3076A;G3087T;G3104A;G3118T;T1622C;T1693C;T1745G;T1768G;T1945G;T1978C;T1984A;T2119C;T2165G;T2188C;T2212C;T2255C;T2258C;T2344C;T2501G;T2549C;T2549G;T2579G;T2609G;T3019C;T3094A;T3096G;T3442C;T3455C;T3463A;T917C;T986C; | K535N;D614G;N710D;T734A;N751S;K776N;Q784R;K811R;S813G;M869V;Q895L;Q901R;K1028I;D1041G;E1072G;Q14L;T547I;T573I;T599I;S640F;P665S;A694V;A701V;T761I;R765C;T778N;T778I;T791I;P812S;L841F;A879V;T883I;T1009I;H1159N;P1162S;P322A;L533F;V534I;V534F;E583D;G594V;V615L;G648V;V656F;E661D;G667D;G667A;G669A;Q677H;A701S;G769E;V772F;G799D;L8F;M900I;R1014T;A1015T;A1026T;M1029I;G1035E;V1040F;F541S;F565L;L582R;C590G;C649G;Y660H;C662S;Y707H;V722G;S730P;C738R;L752P;L753S;F782L;I834S;I850T;I850S;V860G;I870S;Y1007H;C1032S;C1032W;F1148L;L1152S;Y1155N;F306S;F329S; |

NTD, N-terminal domain; RBD, receptor-binding domain; S1/S2, protease cleavage site; HR1, heptad repeat 1; CH, central helix; CD, connector domain; HR2, heptad repeat 2; TM, transmembrane domain; CT, cytoplasmic tail; Others, other regions of the spike gene.

**Supplementary Table 4 Information of high-frequent missense mutations shared by at least 15 haplotypes.**

| Domains | Nucleic acid variations | Amino acid variations |
|---|---|---|
| NTD | A130G;A202G;A220G;A362C;A365C;A439C;A443C;A461G;A559C;A606T;A644G;C115T;C145T;C179A;C205T;C227T;C284T;C323A;C368T;C500T;C59T;C673A;C77A;C863T;C882A;C905T;G104T;G124T;G162T;G224C;G224T;G234T;G247T;G265T;G305T;G319T;G388A;G423T;G452T;G456T;G637T;G656T;G682T;G694T;G695C;G711T;G782A;G803T;G868T;G909T;T107G;T140G;T175C;T185G;T190A;T235G;T248C;T269G;T275C;T276G;T317C;T318G;T334C;T363A;T404A;T406G;T422G;T428G;T476G;T47C;T47G;T495G;T530G;T555G;T578C;T580G;T594G;T603A;T686C;T71C;T746G;T772G;T823C; | A123V;A288V;C136G;D198E;D215G;D228Y;D290Y;D294E;E154G;F106L;F106S;F106T;F135Y;F194V;F201L;F275L;F59L;F79V;F92L;F92S;G107C;G219V;G232A;G232C;G261D;G268V;G35V;G75A;G75V;G89C;H49Y;H69Y;I68V;K147Q;K187Q;K202N;L141F;L141W;L229S;L249W;L24S;L303F;L54F;M177R;N121K;N121T;N122T;N148P;N148T;N165K;N185K;N74D;P225T;P26H;P39S;R102I;R237S;R44G;R78S;S112P;S151I;S60Y;T108N;T167I;T20I;T302M;T76I;T95I;V130I;V143G;V159G;V16A;V16G;V193A;V213L;V36G;V42F;V47G;V62G;V83A;V83F;V90G;W152C;W258G;W64R; |
| RBD | A1153C;A1162G;A1217G;A1315G;A1384C;A1386C;C1387A;C1388T;C1495A;C1499A;G1007C;G1037A;G1087A;G1099T;G1144T;G1211T;G1223T;G1279C;G1337T;G1339T;G1426T;G1454T;G1487T;G1505T;T1014G;T1026A;T1041G;T1111G;T1129C;T1174C;T1344G;T1364G;T1392G;T1457G;T1557G; | A363T;C336S;D427H;E406G;F338L;F342L;F347L;F377L;F392L;F464L;F486C;G404V;G446V;G447C;G476C;G485V;G496V;G502V;H519Q;K462N;K462Q;L455W;N388D;N439D;N448K;P463L;P463T;P499T;R346K;R408I;S371A;T385P;T500N;V367F;V382L; |
| S1S2 | C2047T;G2045T;G2054T; | R682L;R683W;R685L; |
| FP | A2470G;T2465C; | L822P;N824D; |
| HR1 | A2734G;A2759C;A2759G;A2760C;A2777C;A2858C;A2861C;A2892C;A2905C;A2905G;A2906T;A2931T;C2812T;C2873A;C2873T;C2903T;G2824T;G2837T;G2872T;G2911T;G2912T;T2732C | A942S;A958D;A958S;A958V;F927I;F927S;F970L;F970W;G946V;G971C;G971V;K964N;L938F;L977F;N953T;N969D;N969H;N969I;N969K;Q920H;Q920P;Q920R;Q926P;Q954P;S929R;S968F;T912A;V911A; |

| | | |
|---|---|---|
| | ;T2779A;T2787G;T2907A;T2910G; | |
| CH | G2995T;T3002G; | G999C;L1001R; |
| CD | A3292G;A3332G;A3338G;A3355G;A3400C;A3413T;C3247T;C3407T;G3278C;G3296A;G3306T;G3320T;G3371T;G3392T;T3266G;T3283C;T3284C;T3284G;T3307G;T3309G;T3328C;T3361C;T3363G;T3410G;T3412A;T3422G; | E1111G;F1089C;F1095C;F1095L;F1095S;F1103L;F1103V;F1121L;G1093A;G1099D;G1124V;G1131V;H1083Y;L1141W;N1098D;N1119D;N1134H;Q1113R;R1107M;T1136I;V1137G;W1102C;Y1110H;Y1138F;Y1138N; |
| HR2 | A3539C;A3540C;A3541C;A3545G;A3571G;A3575T;A3584C;G3487T;G3512A;T3536C; | D1163Y;E1182G;E1195A;G1171D;I1179T;K1181Q;K1191E;N1192I;Q1180H;Q1180P; |
| TM&CT | A3632C;A3763G;G3650T;G3655T;G3711C;G3761A;G3769T;G3790T;T3640C;T3643C;T3647C;T3703G; | C1235G;C1254Y;D1257Y;G1219C;I1216T;K1211T;K1255E;M1237I;V1264L;W1214R;W1217L;Y1215H; |
| Others | A1582G;A1586G;A1618C;A1669C;A1670C;A1673G;A1841G;A1969G;A1990C;A2008G;A2105G;A2321C;A2324G;A2326C;A2328T;A2330C;A2330T;A2351G;A2401T;A2411C;A2432G;A2437G;A2507C;A2605G;A3083T;A3143G;A3215G;A3216C;A3460G;A3617C;A41T;A926G;C1715T;C1718T;C1862A;C1891T;C1892T;C1919T;C1993A;C1993T;C2102T;C2143A;C2182A;C2282T;C2293T;C2333A;C2372T;C2377A;C2408T;C2434A;C2521T;C2545A;C25T;C2636T;C2648T;C2689T;C3026T;C3156A;C3170A;C3170T;C3206A;C3475A;C3485T;C968T;G1599T;G1600T;G1634T;G1674T;G1740T;G1781T;G1843C;G1898T;G1943T;G1957T;G1966T;G2000A;G2000C;G2006C;G2066C;G2101T;G2306A;G2314T;G2392T;G2395T;G2396T;G24C;G2513A;G2569T;G2578T;G2700T;G3041C;G3043A;G3087T;G3118A;G3118T;G3137T;G3208T;G3447T;G932T;T1571G;T1598G;T1693C;T1722G;T1745G; | K528E;K529R;N540H;K557P;K557Q;K557P;K557T;K558G;K558R;D614G;N657D;N657G;I664L;I670V;E702G;Q774P;D775G;K776L;K776Q;K776N;N777T;N777I;Q784R;N801Y;Q804P;K811R;S813G;Q836P;M869V;K1028I;H1048R;E1072G;E1072D;E1072Y;K1154E;Y1206S;Q14L;E309G;T572I;T573I;P621H;P631F;P631S;P631F;P631L;S640F;P665T;P665S;A701V;P715T;P728T;T761I;R765C;T778N;T791I;P793T;S803F;S803L;P812T;L841F;L849I;P9S;A879V;T883I;P897S;T1009I;F1052L;P1057H;P1057L;P1069H;H1159N;P1162L;T323I;L533F;V534F;G545V;K558N;Q580H;G594V;V615L;W633L;G648V;A653S;V656F;G667D;G667A;G667P;G669A;S689T;A701S;G769E;V772F;G798C;G799C;G799V;L8F;G838D;G857C;V860F;M900I;R1014T;A1015T;M1029I;V1040I;V1040F;G1046V;A1070S;K1149N;G311V;V524G;L533W;F565L;D574E;L582R;C590G;F592L;V6G;V635G;C649G;Y660H;C662S;Y707H;N717K;V722G;S730P;D745E;L752P;C760G;V781G;F782C;F782W;F782L;F782W;F797L;L8S;F800L;N801K;S803P;I834S;I |

| | | |
|---|---|---|
| | T1768G;T1776G;T17G;T1904G;T1945G;T1978C;T1984A;T2119C;T2151A;T2165G;T2188C;T2235G;T2255C;T2278G;T2342G;T2345G;T2346G;T2391G;T23C;T2400A;T2403A;T2407C;T2501G;T2549G;T2579G;T2582G;T2609G;T3019C;T3094A;T3094C;T3096G;T3463A;T8G;T986C; | 850S;V860G;L861W;I870S;Y1007H;C1032S;C1032R;C1032W;Y1155N;V3G;F329S; |

NTD, N-terminal domain; RBD, receptor-binding domain; S1/S2, protease cleavage site; FP, fusion peptide; HR1, heptad repeat 1; CH, central helix; CD, connector domain; HR2, heptad repeat 2; TM, transmembrane domain; CT, cytoplasmic tail; Others, other regions of the spike gene.

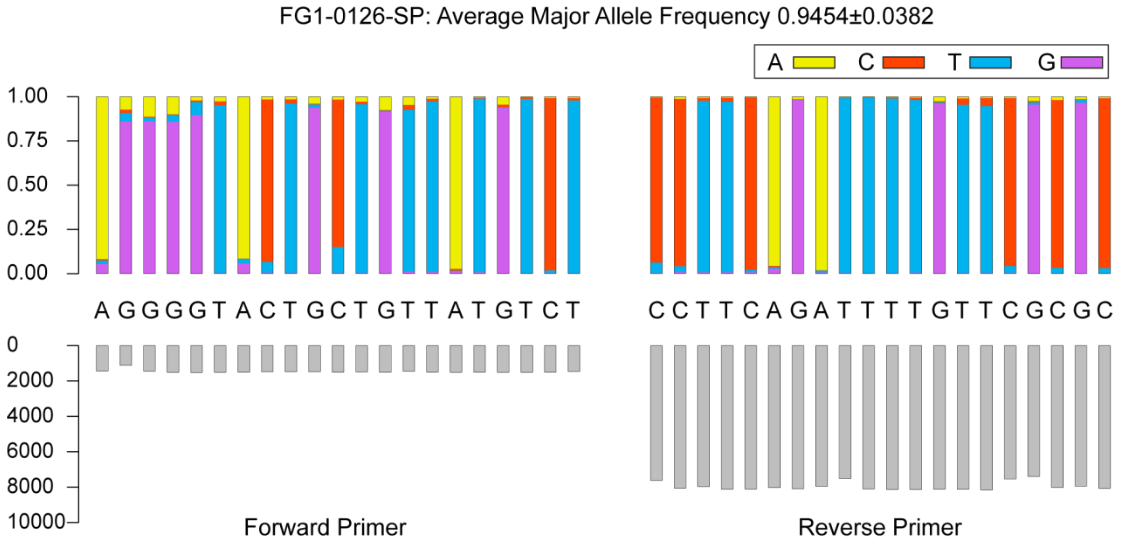
FG1-0126-SP: Average Major Allele Frequency 0.9454±0.0382
A C T G
1.00
0.75
0.50
0.25
0.00
A G G G G T A C T G C T G T T A T G T C T
C C T T C A G A T T T T G T T C G C G C
0
2000
4000
6000
8000
10000
Forward Primer
Reverse Primer

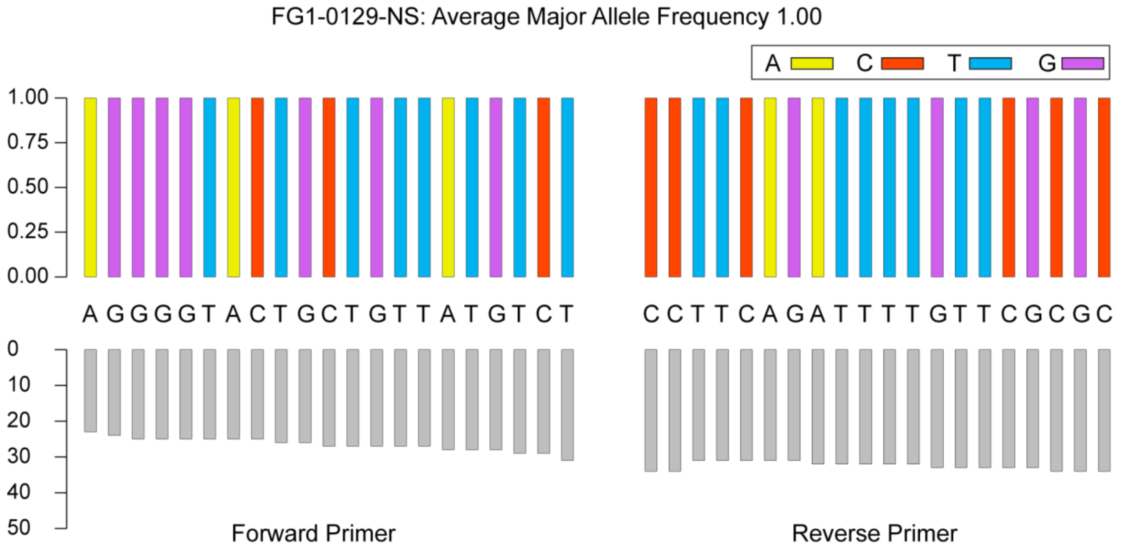
FG1-0129-NS: Average Major Allele Frequency 1.00
A C T G
1.00
0.75
0.50
0.25
0.00
A G G G G T A C T G C T G T T A T G T C T
C C T T C A G A T T T T G T T C G C G C
0
10
20
30
40
50
Forward Primer
Reverse Primer

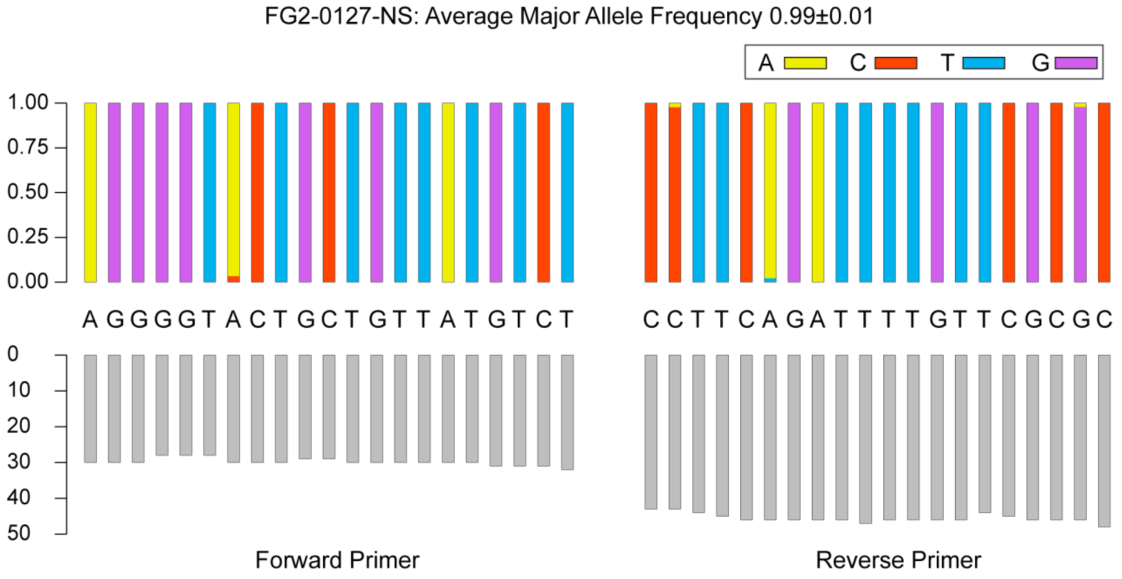
FG2-0127-NS: Average Major Allele Frequency 0.99±0.01
A C T G
1.00
0.75
0.50
0.25
0.00
A G G G G T A C T G C T G T T A T G T C T
C C T T C A G A T T T T G T T C G C G C
0
10
20
30
40
50
Forward Primer
Reverse Primer

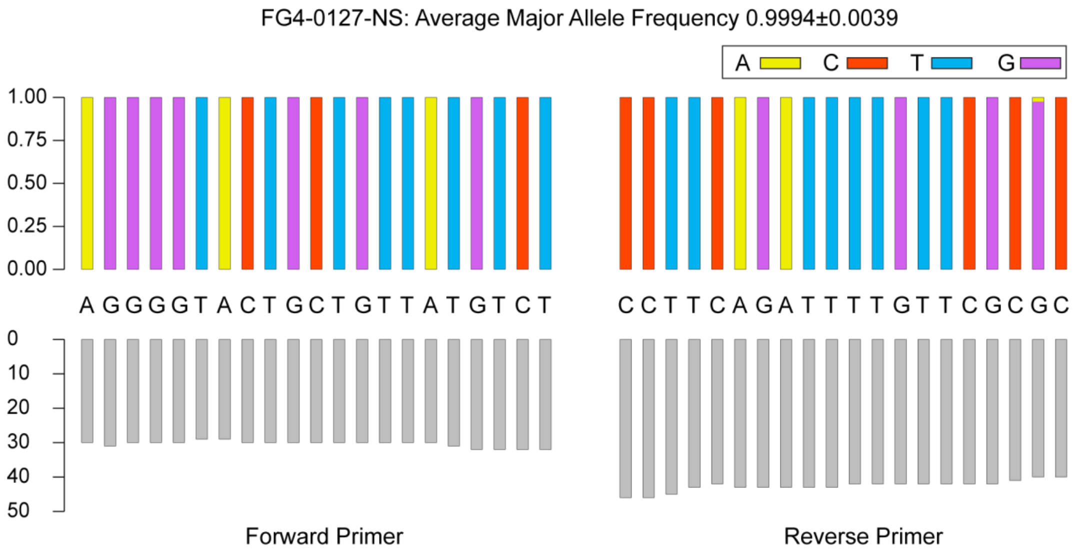
FG4-0127-NS: Average Major Allele Frequency 0.9994±0.0039
A C T G
1.00
0.75
0.50
0.25
0.00
A G G G G T A C T G C T G T T A T G T C T
C C T T C A G A T T T T G T T C G C G C
0
10
20
30
40
50
Forward Primer
Reverse Primer

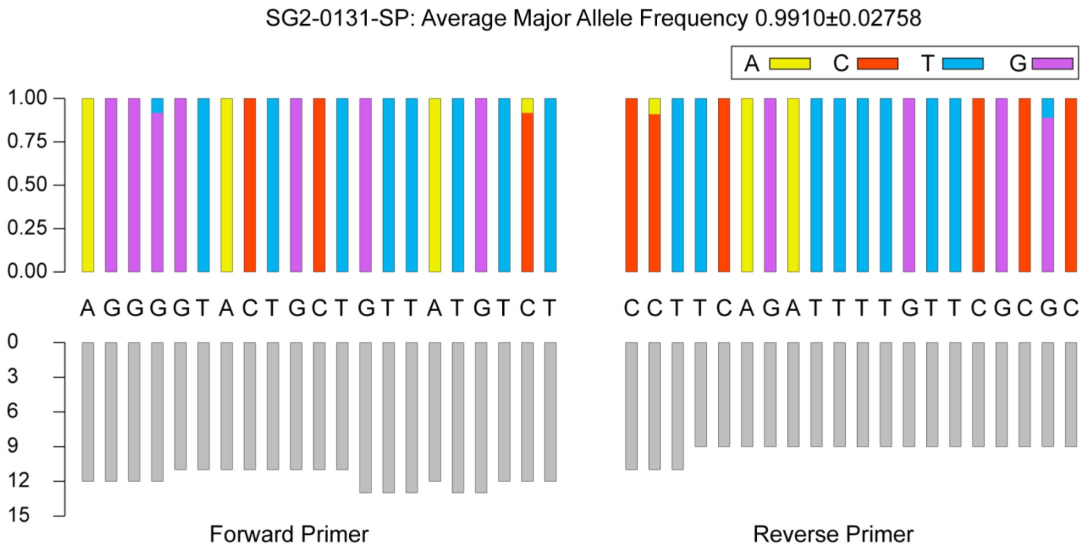
SG2-0131-SP: Average Major Allele Frequency 0.9910±0.02758
A C T G
1.00
0.75
0.50
0.25
0.00
A G G G G T A C T G C T G T T A T G T C T
C C T T C A G A T T T T G T T C G C G C
0
3
6
9
12
15
Forward Primer
Reverse Primer

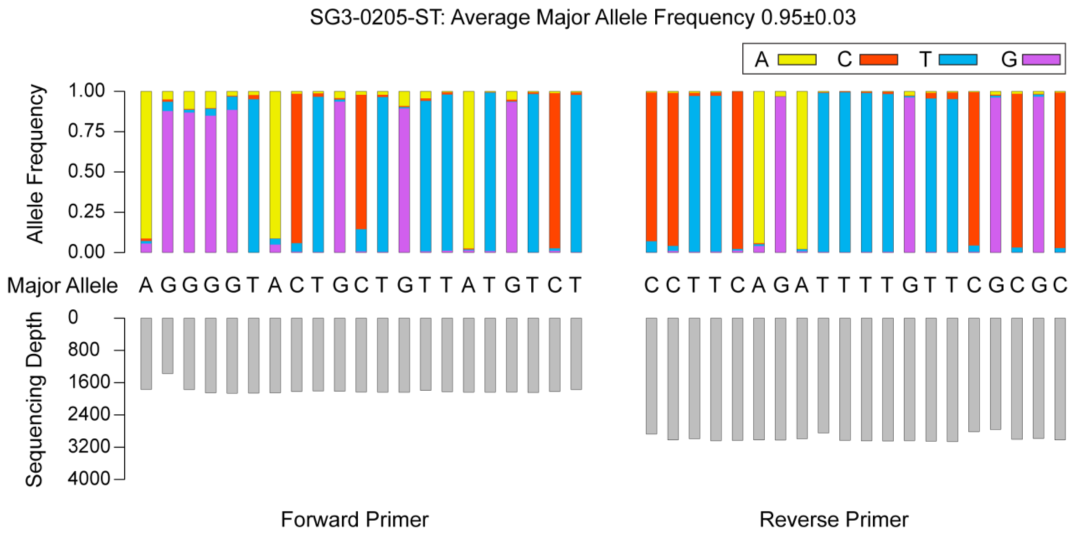
SG3-0205-ST: Average Major Allele Frequency 0.95±0.03
A C T G
Allele Frequency
1.00
0.75
0.50
0.25
0.00
Major Allele
A G G G G T A C T G C T G T T A T G T C T
C C T T C A G A T T T T G T T C G C G C
Sequencing Depth
0
800
1600
2400
3200
4000
Forward Primer
Reverse Primer

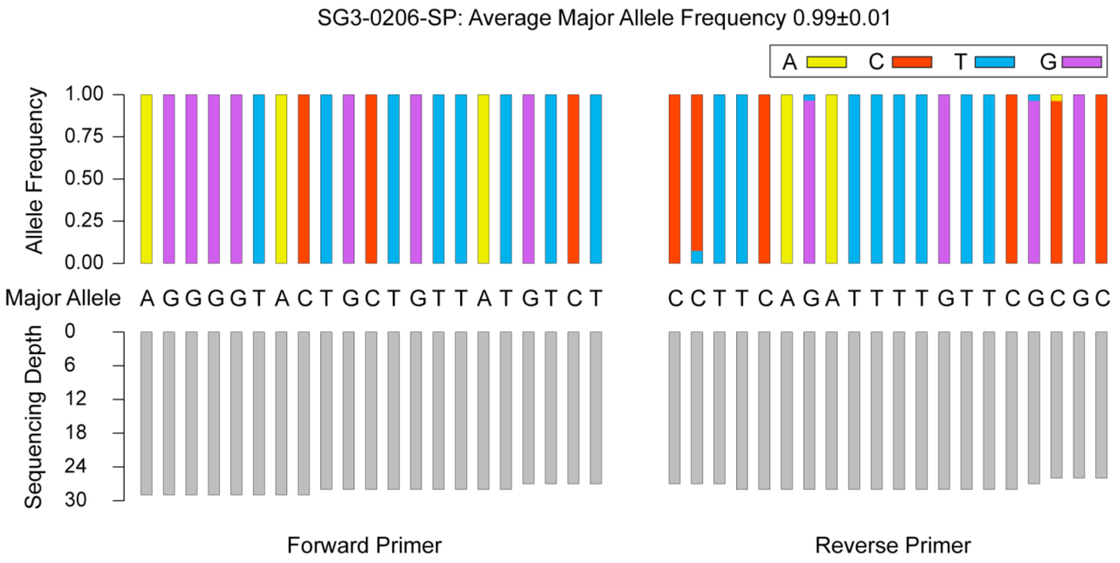


**Supplementary Figure 1 Evaluation of mutation rate of amplified Primers (cDNA).** The upper part shows the frequency distribution of the four bases. The middle represents the major allele. The lower part shows the depth of random NGS sequencing. More than 98% virus RNA contained the identical sequences with the primers of AGGGGTACTGCTGTTATGTCT and CCTTCAGATTTGTTCGCGC.

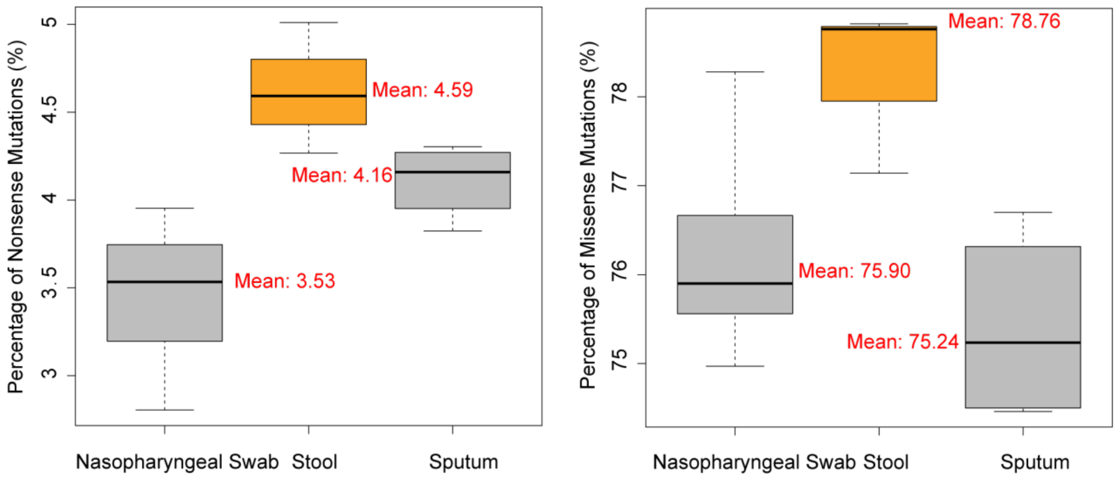


**Supplementary Figure 2 Box plots of the nonsense and missense mutation rate for three sampling types.** Both percentage of nonsense and missense mutation showed obviously higher in samples of stools. The mean values were marked on the figure.

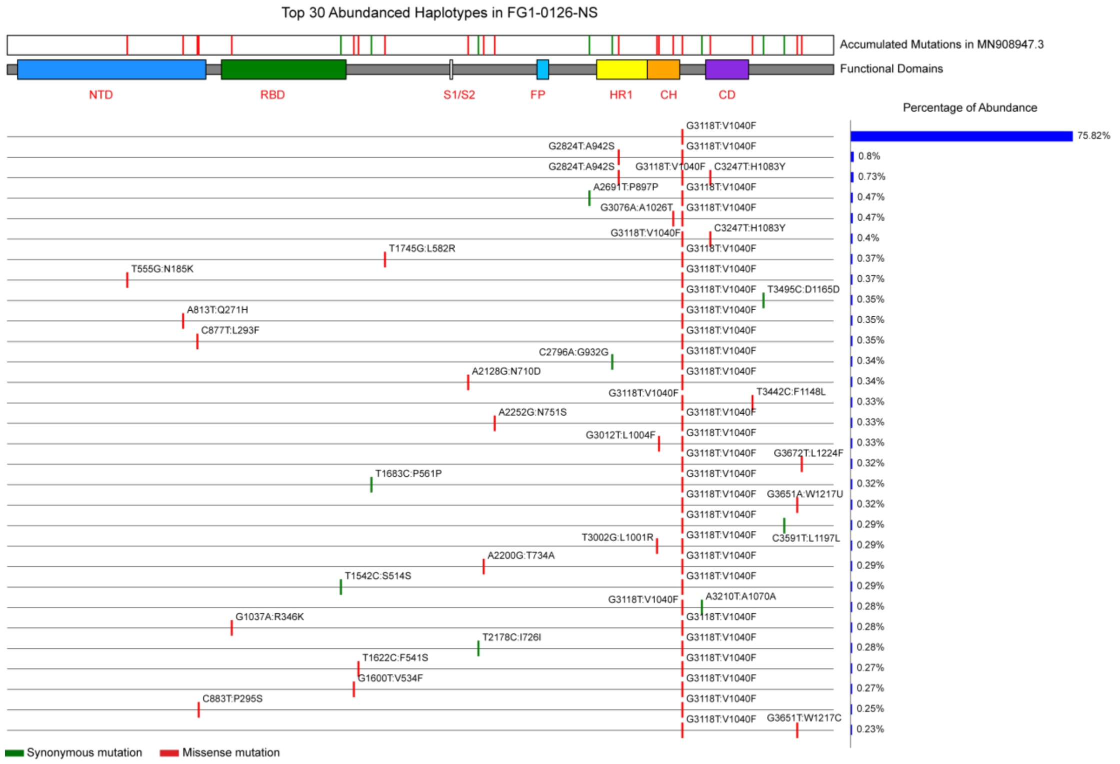
Top 30 Abundanced Haplotypes in FG1-0126-NS
Accumulated Mutations in MN908947.3
Functional Domains
NTD
RBD
S1/S2
FP
HR1
CH
CD
Percentage of Abundance
G3118T:V1040F
75.82%
G2824T:A942S
G3118T:V1040F
0.8%
G2824T:A942S
G3118T:V1040F
C3247T:H1083Y
0.73%
A2691T:P897P
G3118T:V1040F
0.47%
G3076A:A1026T
G3118T:V1040F
0.47%
G3118T:V1040F
C3247T:H1083Y
0.4%
T1745G:L582R
G3118T:V1040F
0.37%
T555G:N185K
G3118T:V1040F
0.37%
G3118T:V1040F
T3495C:D1165D
0.35%
A813T:Q271H
G3118T:V1040F
0.35%
C877T:L293F
G3118T:V1040F
0.35%
C2796A:G932G
G3118T:V1040F
0.34%
A2128G:N710D
G3118T:V1040F
0.34%
G3118T:V1040F
T3442C:F1148L
0.33%
A2252G:N751S
G3118T:V1040F
0.33%
G3012T:L1004F
G3118T:V1040F
0.33%
G3118T:V1040F
G3672T:L1224F
0.32%
T1683C:P561P
G3118T:V1040F
0.32%
G3118T:V1040F
G3651A:W1217U
0.32%
G3118T:V1040F
0.29%
T3002G:L1001R
G3118T:V1040F
C3591T:L1197L
0.29%
A2200G:T734A
G3118T:V1040F
0.29%
T1542C:S514S
G3118T:V1040F
0.29%
G3118T:V1040F
A3210T:A1070A
0.28%
G1037A:R346K
G3118T:V1040F
0.28%
T2178C:I726I
G3118T:V1040F
0.28%
T1622C:F541S
G3118T:V1040F
0.27%
G1600T:V534F
G3118T:V1040F
0.27%
C883T:P295S
G3118T:V1040F
0.25%
G3118T:V1040F
G3651T:W1217C
0.23%
Synonymous mutation
Missense mutation

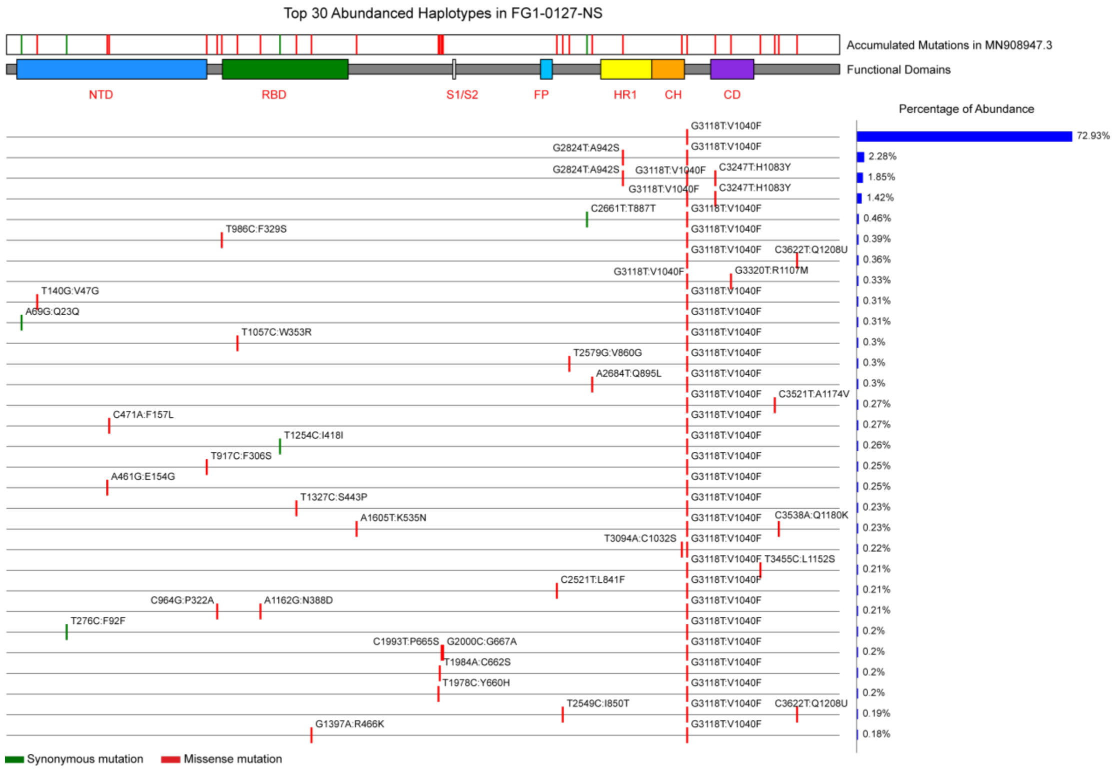
Top 30 Abundanced Haplotypes in FG1-0127-NS
Accumulated Mutations in MN908947.3
Functional Domains
NTD
RBD
S1/S2
FP
HR1
CH
CD
Percentage of Abundance
G3118T:V1040F
72.93%
G2824T:A942S
G3118T:V1040F
2.28%
G2824T:A942S
G3118T:V1040F
C3247T:H1083Y
1.85%
G3118T:V1040F
C3247T:H1083Y
1.42%
C2661T:T887T
G3118T:V1040F
0.46%
T986C:F329S
G3118T:V1040F
0.39%
G3118T:V1040F
C3622T:Q1208U
0.36%
G3118T:V1040F
G3320T:R1107M
0.33%
T140G:V47G
G3118T:V1040F
0.31%
A69G:Q23Q
G3118T:V1040F
0.31%
T1057C:W353R
G3118T:V1040F
0.3%
T2579G:V860G
G3118T:V1040F
0.3%
A2684T:Q895L
G3118T:V1040F
0.3%
G3118T:V1040F
C3521T:A1174V
0.27%
C471A:F157L
G3118T:V1040F
0.27%
T1254C:I418I
G3118T:V1040F
0.26%
T917C:F306S
G3118T:V1040F
0.25%
A461G:E154G
G3118T:V1040F
0.25%
T1327C:S443P
G3118T:V1040F
0.23%
A1605T:K535N
G3118T:V1040F
C3538A:Q1180K
0.23%
T3094A:C1032S
G3118T:V1040F
0.22%
G3118T:V1040F
T3455C:L1152S
0.21%
C2521T:L841F
G3118T:V1040F
0.21%
C964G:P322A
A1162G:N388D
G3118T:V1040F
0.21%
T276C:F92F
G3118T:V1040F
0.2%
C1993T:P665S
G2000C:G667A
G3118T:V1040F
0.2%
T1984A:C662S
G3118T:V1040F
0.2%
T1978C:Y660H
G3118T:V1040F
0.2%
T2549C:I850T
G3118T:V1040F
C3622T:Q1208U
0.19%
G1397A:R466K
G3118T:V1040F
0.18%
Synonymous mutation
Missense mutation

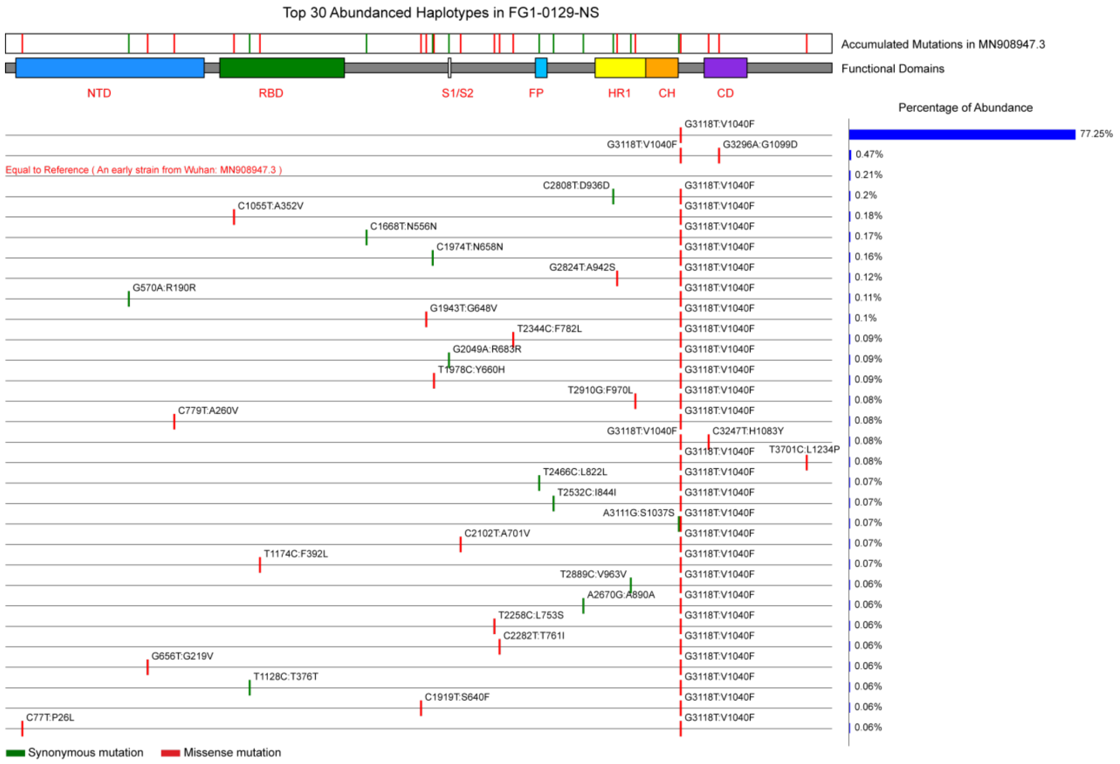
Top 30 Abundanced Haplotypes in FG1-0129-NS
Accumulated Mutations in MN908947.3
Functional Domains
NTD
RBD
S1/S2
FP
HR1
CH
CD
Percentage of Abundance
G3118T:V1040F
77.25%
G3118T:V1040F
G3296A:G1099D
0.47%
Equal to Reference ( An early strain from Wuhan: MN908947.3 )
0.21%
C2808T:D936D
G3118T:V1040F
0.2%
C1055T:A352V
G3118T:V1040F
0.18%
C1668T:N556N
G3118T:V1040F
0.17%
C1974T:N658N
G3118T:V1040F
0.16%
G2824T:A942S
G3118T:V1040F
0.12%
G570A:R190R
G3118T:V1040F
0.11%
G1943T:G648V
G3118T:V1040F
0.1%
T2344C:F782L
G3118T:V1040F
0.09%
G2049A:R683R
G3118T:V1040F
0.09%
T1978C:Y660H
G3118T:V1040F
0.09%
T2910G:F970L
G3118T:V1040F
0.08%
C779T:A260V
G3118T:V1040F
0.08%
G3118T:V1040F
C3247T:H1083Y
0.08%
G3118T:V1040F
T3701C:L1234P
0.08%
T2466C:L822L
G3118T:V1040F
0.07%
T2532C:I844I
G3118T:V1040F
0.07%
A3111G:S1037S
G3118T:V1040F
0.07%
C2102T:A701V
G3118T:V1040F
0.07%
T1174C:F392L
G3118T:V1040F
0.07%
T2889C:V963V
G3118T:V1040F
0.06%
A2670G:A890A
G3118T:V1040F
0.06%
T2258C:L753S
G3118T:V1040F
0.06%
C2282T:T761I
G3118T:V1040F
0.06%
G656T:G219V
G3118T:V1040F
0.06%
T1128C:T376T
G3118T:V1040F
0.06%
C1919T:S640F
G3118T:V1040F
0.06%
C77T:P26L
G3118T:V1040F
0.06%
Synonymous mutation
Missense mutation

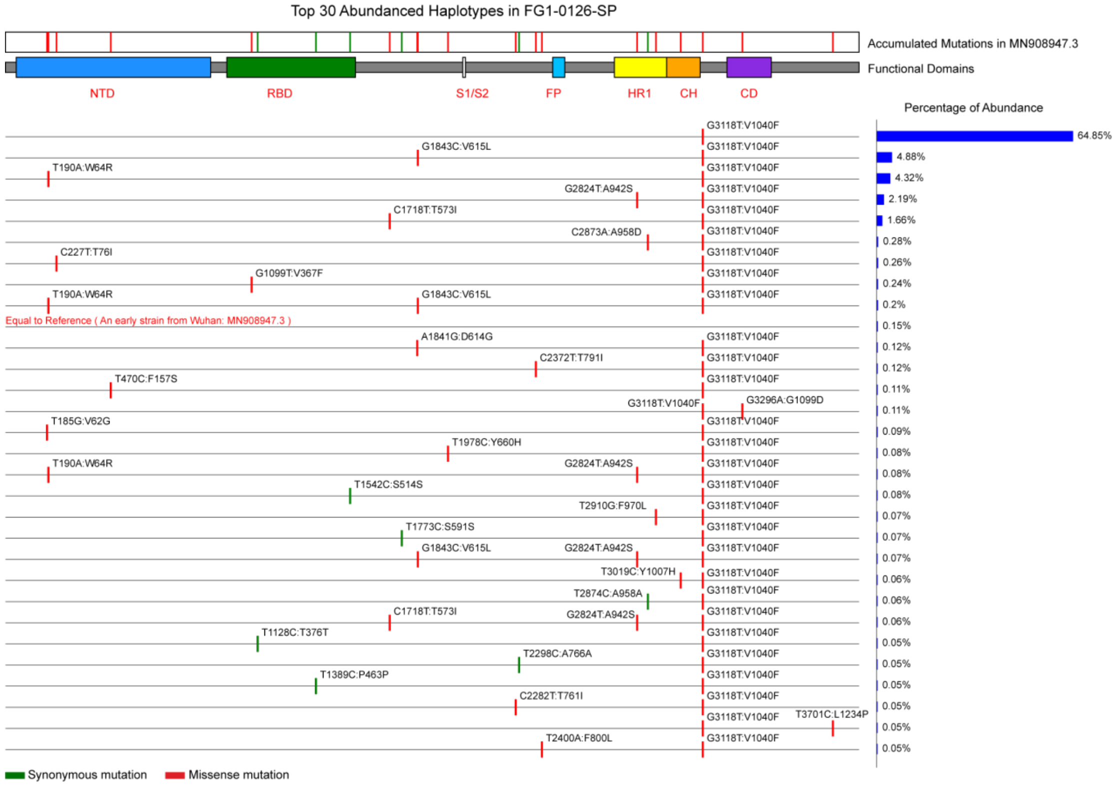
Top 30 Abundanced Haplotypes in FG1-0126-SP
Accumulated Mutations in MN908947.3
Functional Domains
NTD
RBD
S1/S2
FP
HR1
CH
CD
Percentage of Abundance
G3118T:V1040F
64.85%
G1843C:V615L
G3118T:V1040F
4.88%
T190A:W64R
G3118T:V1040F
4.32%
G2824T:A942S
G3118T:V1040F
2.19%
C1718T:T573I
G3118T:V1040F
1.66%
C2873A:A958D
G3118T:V1040F
0.28%
C227T:T76I
G3118T:V1040F
0.26%
G1099T:V367F
G3118T:V1040F
0.24%
T190A:W64R
G1843C:V615L
G3118T:V1040F
0.2%
Equal to Reference ( An early strain from Wuhan: MN908947.3 )
0.15%
A1841G:D614G
G3118T:V1040F
0.12%
C2372T:T791I
G3118T:V1040F
0.12%
T470C:F157S
G3118T:V1040F
0.11%
G3118T:V1040F
G3296A:G1099D
0.11%
T185G:V62G
G3118T:V1040F
0.09%
T1978C:Y660H
G3118T:V1040F
0.08%
T190A:W64R
G2824T:A942S
G3118T:V1040F
0.08%
T1542C:S514S
G3118T:V1040F
0.08%
T2910G:F970L
G3118T:V1040F
0.07%
T1773C:S591S
G3118T:V1040F
0.07%
G1843C:V615L
G2824T:A942S
G3118T:V1040F
0.07%
T3019C:Y1007H
G3118T:V1040F
0.06%
T2874C:A958A
G3118T:V1040F
0.06%
C1718T:T573I
G2824T:A942S
G3118T:V1040F
0.06%
T1128C:T376T
G3118T:V1040F
0.05%
T2298C:A766A
G3118T:V1040F
0.05%
T1389C:P463P
G3118T:V1040F
0.05%
C2282T:T761I
G3118T:V1040F
0.05%
G3118T:V1040F
T3701C:L1234P
0.05%
T2400A:F800L
G3118T:V1040F
0.05%
Synonymous mutation
Missense mutation

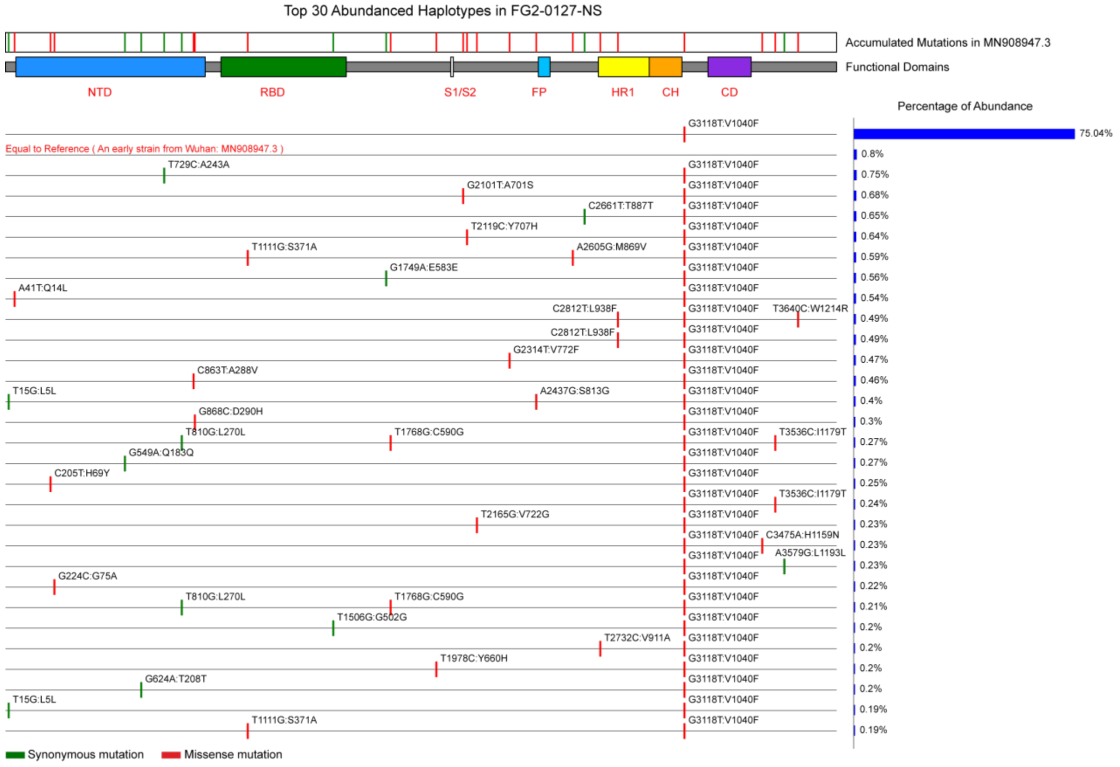

Top 30 Abundanced Haplotypes in FG2-0127-NS
Accumulated Mutations in MN908947.3
Functional Domains
NTD
RBD
S1/S2
FP
HR1
CH
CD
Percentage of Abundance
Equal to Reference ( An early strain from Wuhan: MN908947.3 )
G3118T:V1040F
75.04%
Synonymous mutation
Missense mutation


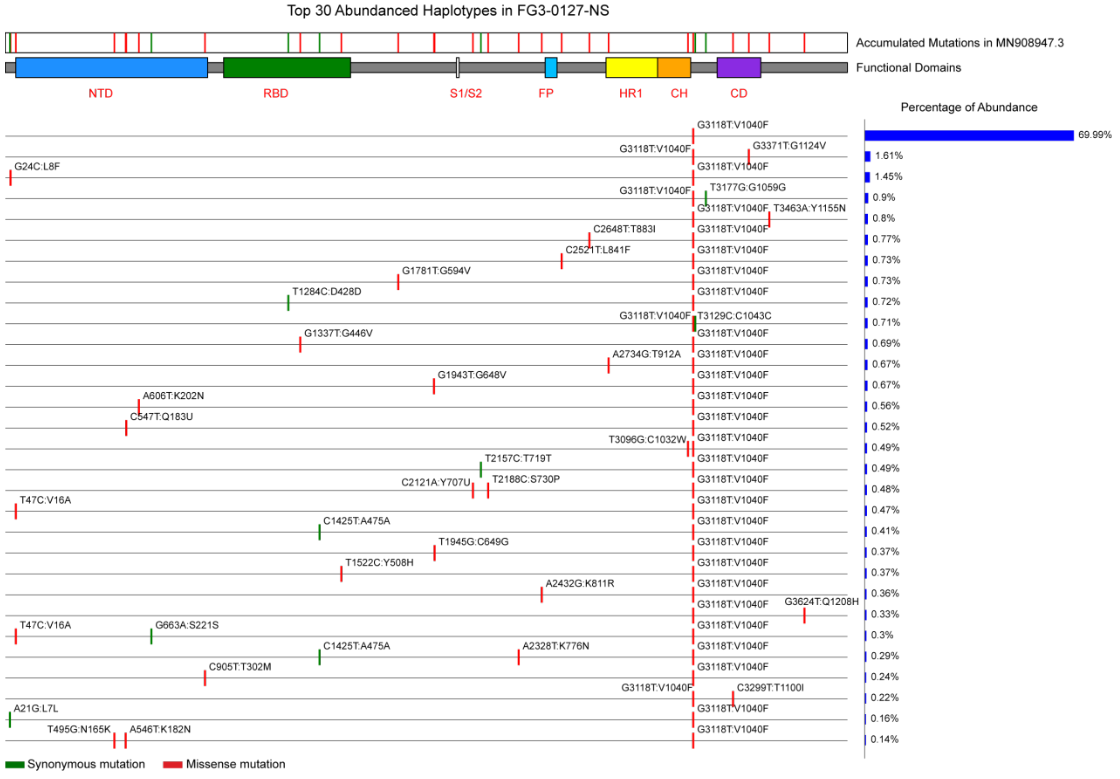

Top 30 Abundanced Haplotypes in FG3-0127-NS
Accumulated Mutations in MN908947.3
Functional Domains
NTD
RBD
S1/S2
FP
HR1
CH
CD
Percentage of Abundance
G3118T:V1040F
69.99%
Synonymous mutation
Missense mutation

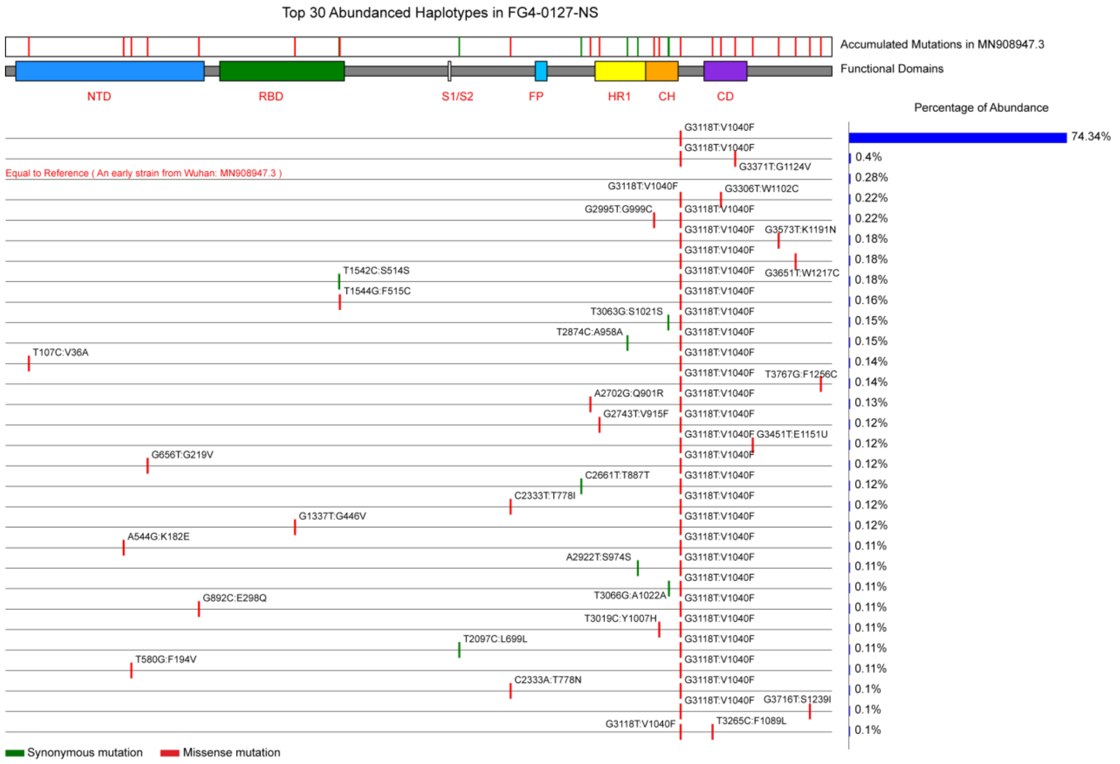
Top 30 Abundanced Haplotypes in FG4-0127-NS
Accumulated Mutations in MN908947.3
Functional Domains
NTD
RBD
S1/S2
FP
HR1
CH
CD
Percentage of Abundance
G3118T:V1040F
74.34%
G3118T:V1040F
G3371T:G1124V
0.4%
Equal to Reference ( An early strain from Wuhan: MN908947.3 )
0.28%
G3118T:V1040F
G3306T:W1102C
0.22%
G2995T:G999C
G3118T:V1040F
0.22%
G3118T:V1040F
G3573T:K1191N
0.18%
G3118T:V1040F
0.18%
T1542C:S514S
G3118T:V1040F
G3651T:W1217C
0.18%
T1544G:F515C
G3118T:V1040F
0.16%
T3063G:S1021S
G3118T:V1040F
0.15%
T2874C:A958A
G3118T:V1040F
0.15%
T107C:V36A
G3118T:V1040F
0.14%
G3118T:V1040F
T3767G:F1256C
0.14%
A2702G:Q901R
G3118T:V1040F
0.13%
G2743T:V915F
G3118T:V1040F
0.12%
G3118T:V1040F
G3451T:E1151U
0.12%
G656T:G219V
G3118T:V1040F
0.12%
C2661T:T887T
G3118T:V1040F
0.12%
C2333T:T778I
G3118T:V1040F
0.12%
G1337T:G446V
G3118T:V1040F
0.12%
A544G:K182E
G3118T:V1040F
0.11%
A2922T:S974S
G3118T:V1040F
0.11%
G3118T:V1040F
0.11%
G892C:E298Q
T3066G:A1022A
G3118T:V1040F
0.11%
T3019C:Y1007H
G3118T:V1040F
0.11%
T2097C:L699L
G3118T:V1040F
0.11%
T580G:F194V
G3118T:V1040F
0.11%
C2333A:T778N
G3118T:V1040F
0.1%
G3118T:V1040F
G3716T:S1239I
0.1%
G3118T:V1040F
T3265C:F1089L
0.1%
Synonymous mutation
Missense mutation

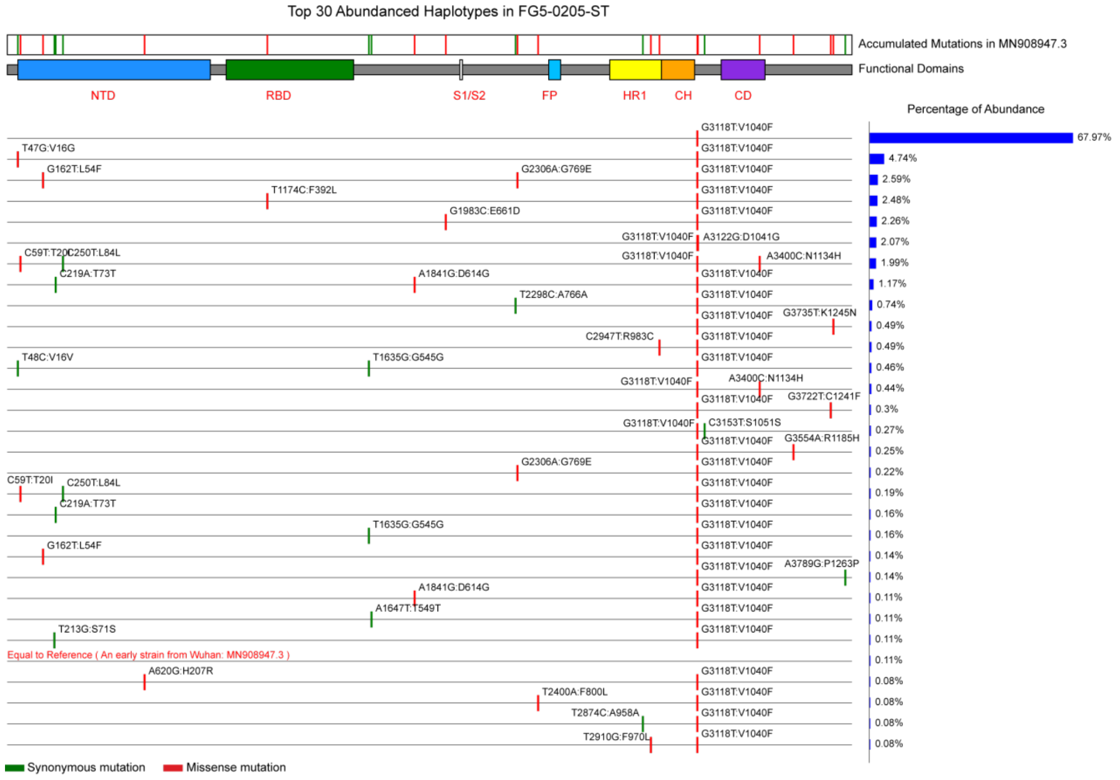
Top 30 Abundanced Haplotypes in FG5-0205-ST
Accumulated Mutations in MN908947.3
Functional Domains
NTD
RBD
S1/S2
FP
HR1
CH
CD
Percentage of Abundance
G3118T:V1040F
67.97%
T47G:V16G
G3118T:V1040F
4.74%
G162T:L54F
G2306A:G769E
G3118T:V1040F
2.59%
T1174C:F392L
G3118T:V1040F
2.48%
G1983C:E661D
G3118T:V1040F
2.26%
G3118T:V1040F
A3122G:D1041G
2.07%
C59T:T20I
C250T:L84L
G3118T:V1040F
A3400C:N1134H
1.99%
C219A:T73T
A1841G:D614G
G3118T:V1040F
1.17%
T2298C:A766A
G3118T:V1040F
0.74%
G3118T:V1040F
G3735T:K1245N
0.49%
C2947T:R983C
G3118T:V1040F
0.49%
T48C:V16V
T1635G:G545G
G3118T:V1040F
0.46%
G3118T:V1040F
A3400C:N1134H
0.44%
G3118T:V1040F
G3722T:C1241F
0.3%
G3118T:V1040F
C3153T:S1051S
0.27%
G3118T:V1040F
G3554A:R1185H
0.25%
G2306A:G769E
G3118T:V1040F
0.22%
C59T:T20I
C250T:L84L
G3118T:V1040F
0.19%
C219A:T73T
G3118T:V1040F
0.16%
T1635G:G545G
G3118T:V1040F
0.16%
G162T:L54F
G3118T:V1040F
0.14%
G3118T:V1040F
A3789G:P1263P
0.14%
A1841G:D614G
G3118T:V1040F
0.11%
A1647T:T549T
G3118T:V1040F
0.11%
T213G:S71S
G3118T:V1040F
0.11%
Equal to Reference ( An early strain from Wuhan: MN908947.3 )
0.11%
A620G:H207R
G3118T:V1040F
0.08%
T2400A:F800L
G3118T:V1040F
0.08%
T2874C:A958A
G3118T:V1040F
0.08%
T2910G:F970L
G3118T:V1040F
0.08%
Synonymous mutation
Missense mutation

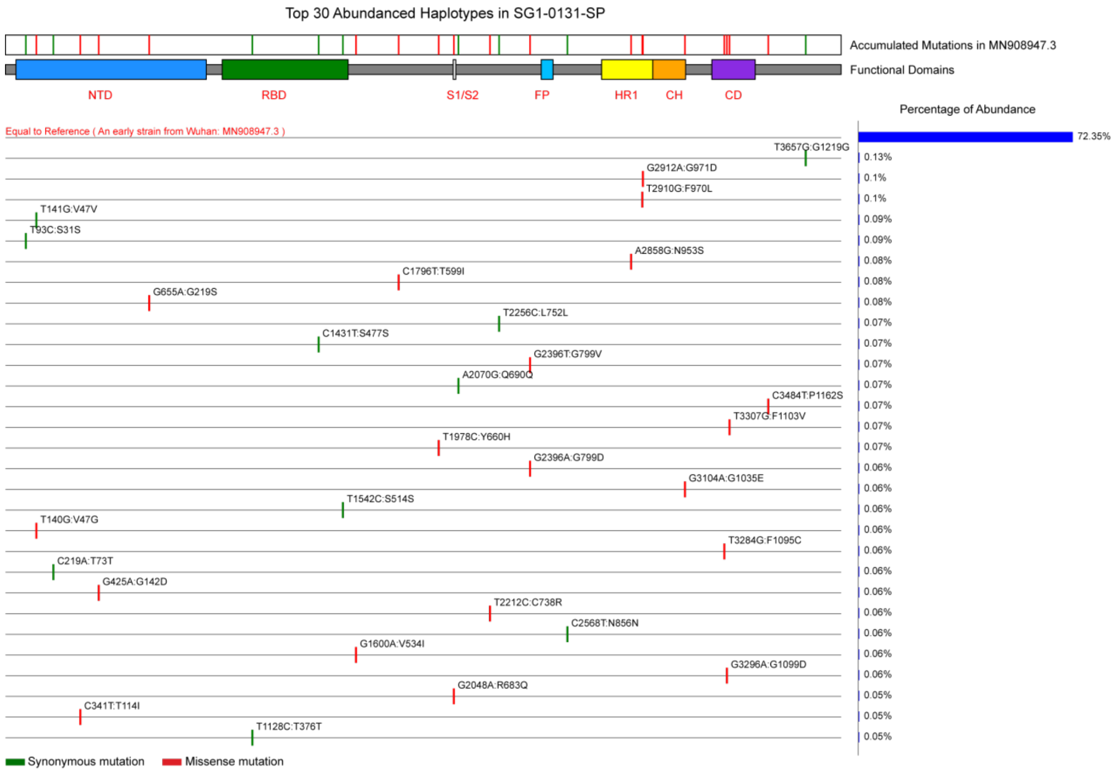
Top 30 Abundanced Haplotypes in SG1-0131-SP
Accumulated Mutations in MN908947.3
Functional Domains
NTD
RBD
S1/S2
FP
HR1
CH
CD
Percentage of Abundance
Equal to Reference ( An early strain from Wuhan: MN908947.3 )
72.35%
T3657G:G1219G
0.13%
G2912A:G971D
0.1%
T2910G:F970L
0.1%
T141G:V47V
0.09%
T93C:S31S
0.09%
A2858G:N953S
0.08%
C1796T:T599I
0.08%
G655A:G219S
0.08%
T2256C:L752L
0.07%
C1431T:S477S
0.07%
G2396T:G799V
0.07%
A2070G:Q690Q
0.07%
C3484T:P1162S
0.07%
T3307G:F1103V
0.07%
T1978C:Y660H
0.07%
G2396A:G799D
0.06%
G3104A:G1035E
0.06%
T1542C:S514S
0.06%
T140G:V47G
0.06%
T3284G:F1095C
0.06%
C219A:T73T
0.06%
G425A:G142D
0.06%
T2212C:C738R
0.06%
C2568T:N856N
0.06%
G1600A:V534I
0.06%
G3296A:G1099D
0.06%
G2048A:R683Q
0.05%
C341T:T114I
0.05%
T1128C:T376T
0.05%
Synonymous mutation
Missense mutation

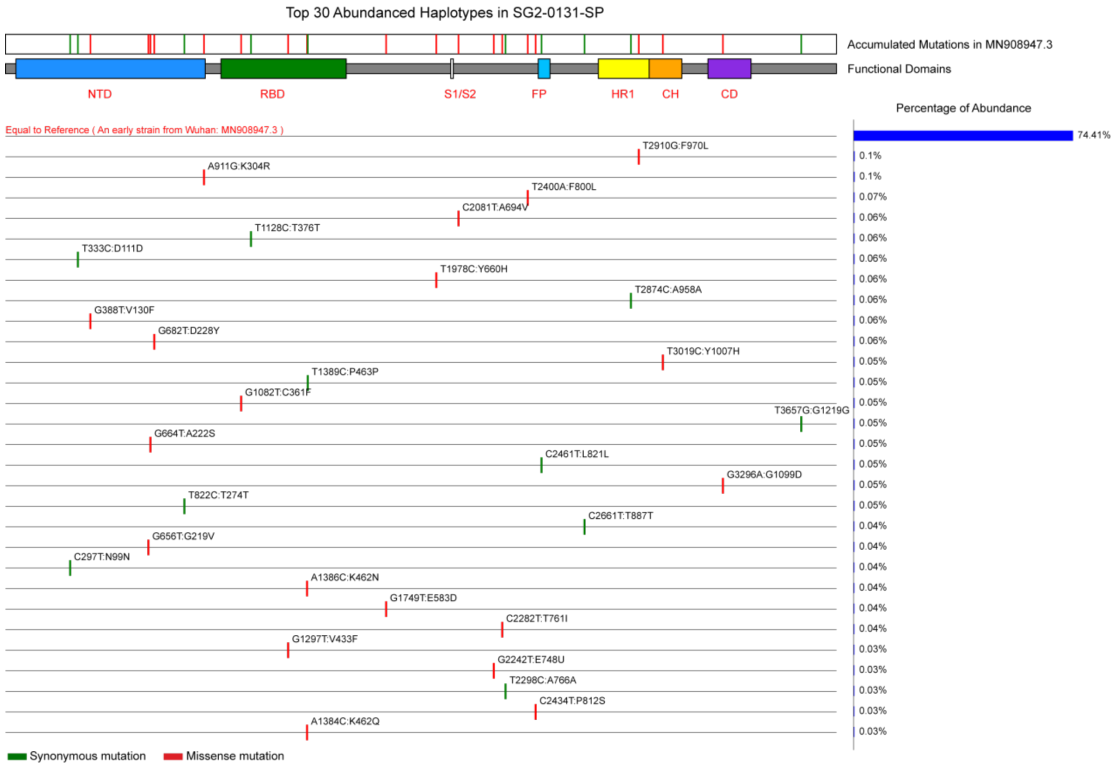
Top 30 Abundanced Haplotypes in SG2-0131-SP
Accumulated Mutations in MN908947.3
Functional Domains
NTD
RBD
S1/S2
FP
HR1
CH
CD
Percentage of Abundance
Equal to Reference ( An early strain from Wuhan: MN908947.3 )
74.41%
T2910G:F970L
0.1%
A911G:K304R
0.1%
T2400A:F800L
0.07%
C2081T:A694V
0.06%
T1128C:T376T
0.06%
T333C:D111D
0.06%
T1978C:Y660H
0.06%
T2874C:A958A
0.06%
G388T:V130F
0.06%
G682T:D228Y
0.06%
T3019C:Y1007H
0.05%
T1389C:P463P
0.05%
G1082T:C361F
0.05%
T3657G:G1219G
0.05%
G664T:A222S
0.05%
C2461T:L821L
0.05%
G3296A:G1099D
0.05%
T822C:T274T
0.05%
C2661T:T887T
0.04%
G656T:G219V
0.04%
C297T:N99N
0.04%
A1386C:K462N
0.04%
G1749T:E583D
0.04%
C2282T:T761I
0.04%
G1297T:V433F
0.03%
G2242T:E748U
0.03%
T2298C:A766A
0.03%
C2434T:P812S
0.03%
A1384C:K462Q
0.03%
Synonymous mutation
Missense mutation

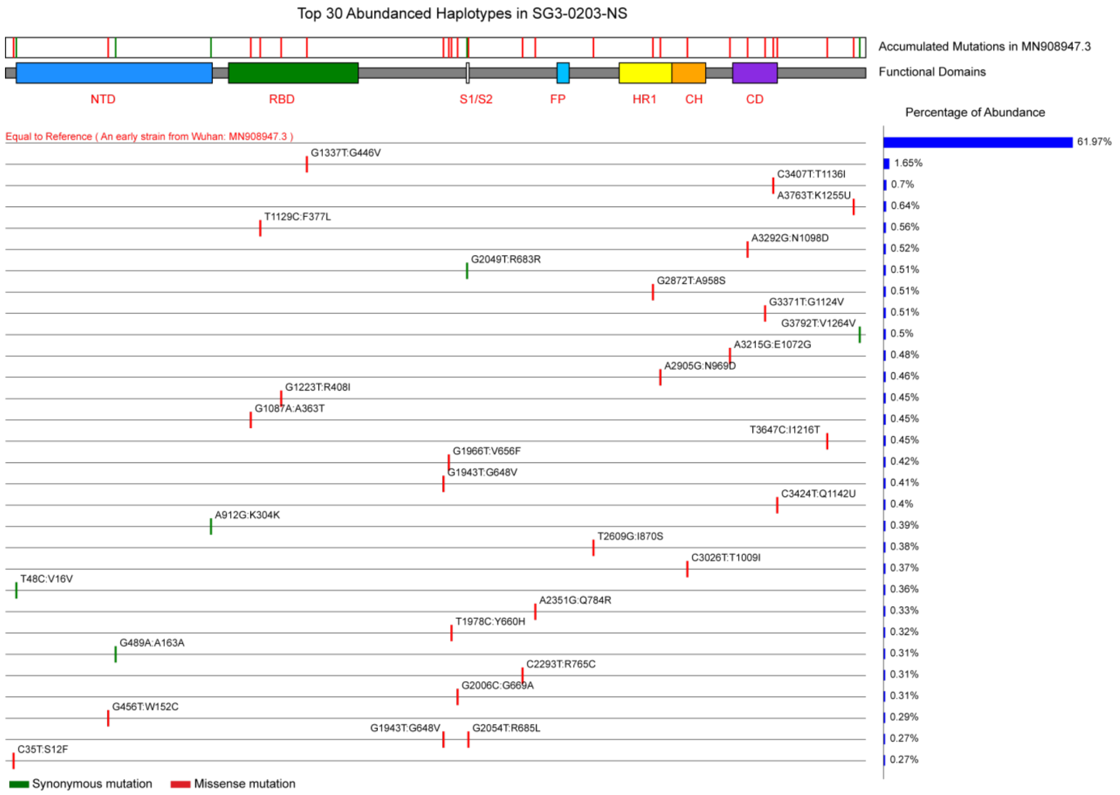
Top 30 Abundanced Haplotypes in SG3-0203-NS
Accumulated Mutations in MN908947.3
Functional Domains
NTD
RBD
S1/S2
FP
HR1
CH
CD
Percentage of Abundance
Equal to Reference ( An early strain from Wuhan: MN908947.3 )
61.97%
G1337T:G446V
1.65%
C3407T:T1136I
0.7%
A3763T:K1255U
0.64%
T1129C:F377L
0.56%
A3292G:N1098D
0.52%
G2049T:R683R
0.51%
G2872T:A958S
0.51%
G3371T:G1124V
0.51%
G3792T:V1264V
0.5%
A3215G:E1072G
0.48%
A2905G:N969D
0.46%
G1223T:R408I
0.45%
G1087A:A363T
0.45%
T3647C:I1216T
0.45%
G1966T:V656F
0.42%
G1943T:G648V
0.41%
C3424T:Q1142U
0.4%
A912G:K304K
0.39%
T2609G:I870S
0.38%
C3026T:T1009I
0.37%
T48C:V16V
0.36%
A2351G:Q784R
0.33%
T1978C:Y660H
0.32%
G489A:A163A
0.31%
C2293T:R765C
0.31%
G2006C:G669A
0.31%
G456T:W152C
0.29%
G1943T:G648V
G2054T:R685L
0.27%
C35T:S12F
0.27%
Synonymous mutation
Missense mutation

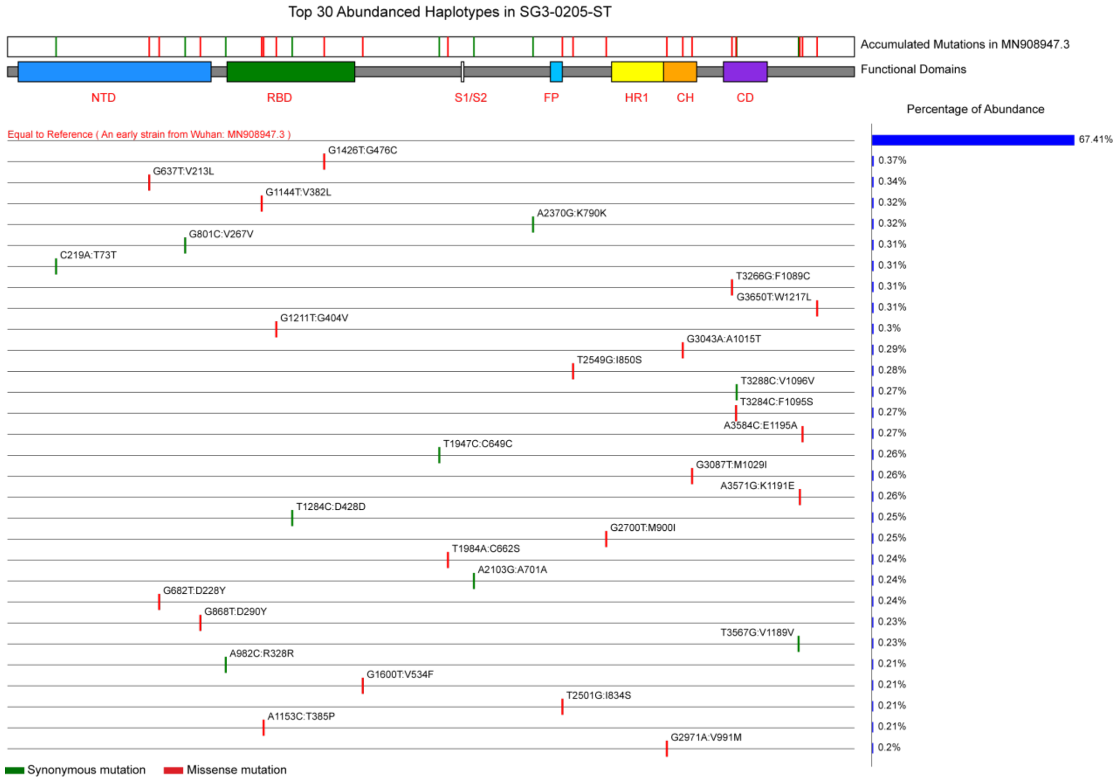
Top 30 Abundanced Haplotypes in SG3-0205-ST
Accumulated Mutations in MN908947.3
Functional Domains
NTD
RBD
S1/S2
FP
HR1
CH
CD
Percentage of Abundance
Equal to Reference ( An early strain from Wuhan: MN908947.3 )
67.41%
G1426T:G476C
0.37%
G637T:V213L
0.34%
G1144T:V382L
0.32%
A2370G:K790K
0.32%
G801C:V267V
0.31%
C219A:T73T
0.31%
T3266G:F1089C
0.31%
G3650T:W1217L
0.31%
G1211T:G404V
0.3%
G3043A:A1015T
0.29%
T2549G:I850S
0.28%
T3288C:V1096V
0.27%
T3284C:F1095S
0.27%
A3584C:E1195A
0.27%
T1947C:C649C
0.26%
G3087T:M1029I
0.26%
A3571G:K1191E
0.26%
T1284C:D428D
0.25%
G2700T:M900I
0.25%
T1984A:C662S
0.24%
A2103G:A701A
0.24%
G682T:D228Y
0.24%
G868T:D290Y
0.23%
T3567G:V1189V
0.23%
A982C:R328R
0.21%
G1600T:V534F
0.21%
T2501G:I834S
0.21%
A1153C:T385P
0.21%
G2971A:V991M
0.2%
Synonymous mutation
Missense mutation

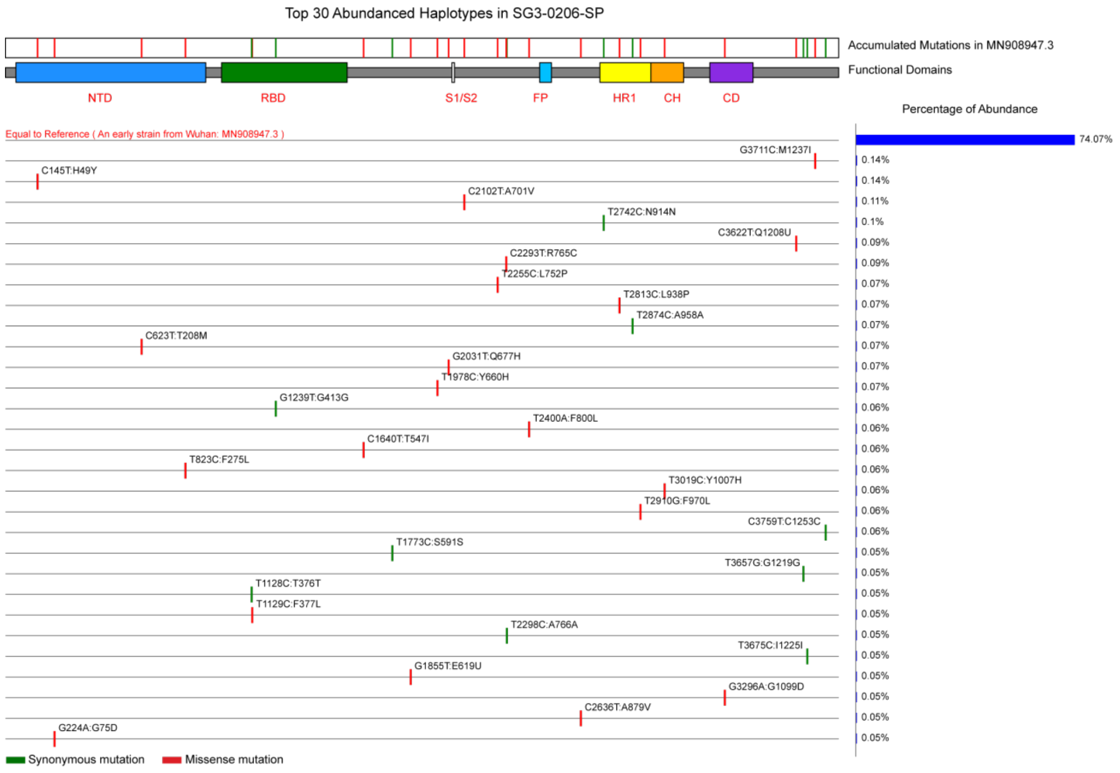


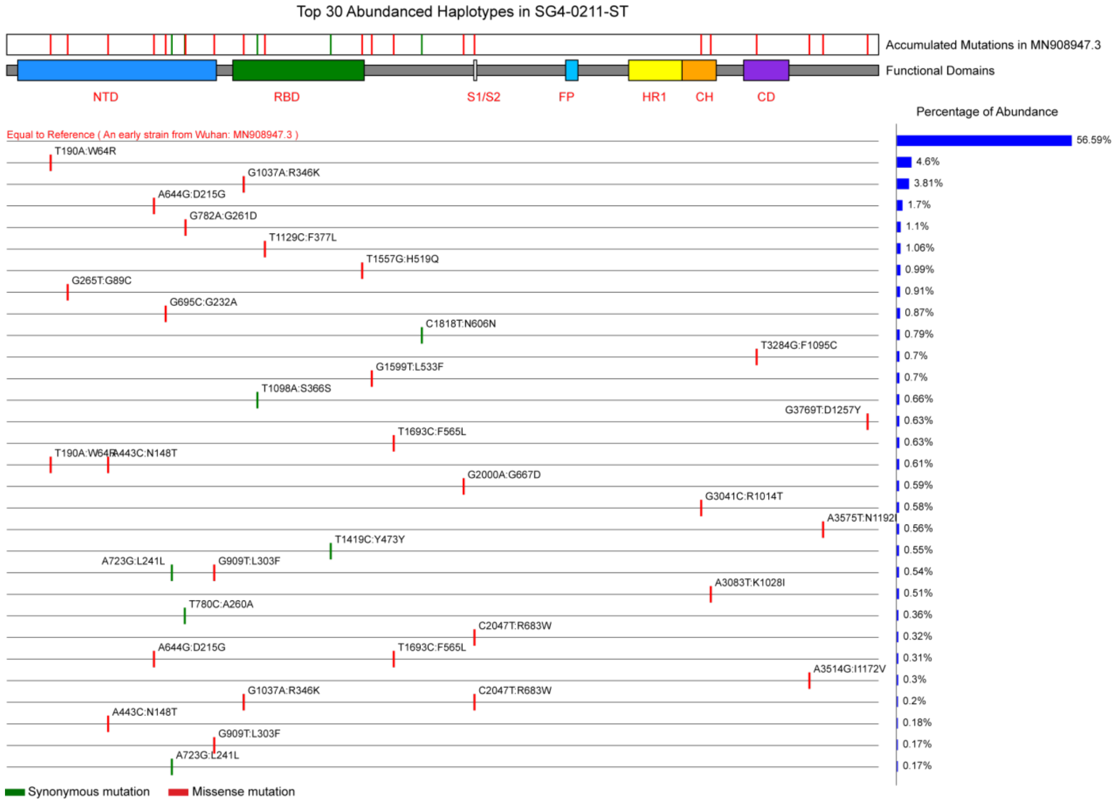


**Supplementary Figure 3 Distribution of top 30 abundant haplotypes of each sample.**

Each line represented a haplotype, green lines were the missense mutations and the red colors were synonymous mutations. The top rectangle represents the cumulative variations from quasispecies haplotypes. Functional domain of spike gene were marked with different colors:

NTD, N-terminal domain; RBD, receptor-binding domain; S1/S2, protease cleavage site; FP, fusion peptide; HR1, heptad repeat 1; CH, central helix; CD, connector domain; HR2, heptad repeat 2; TM, transmembrane domain; CT, cytoplasmic tail. The rightmost bar plots showed the corresponding haplotype abundance (%).

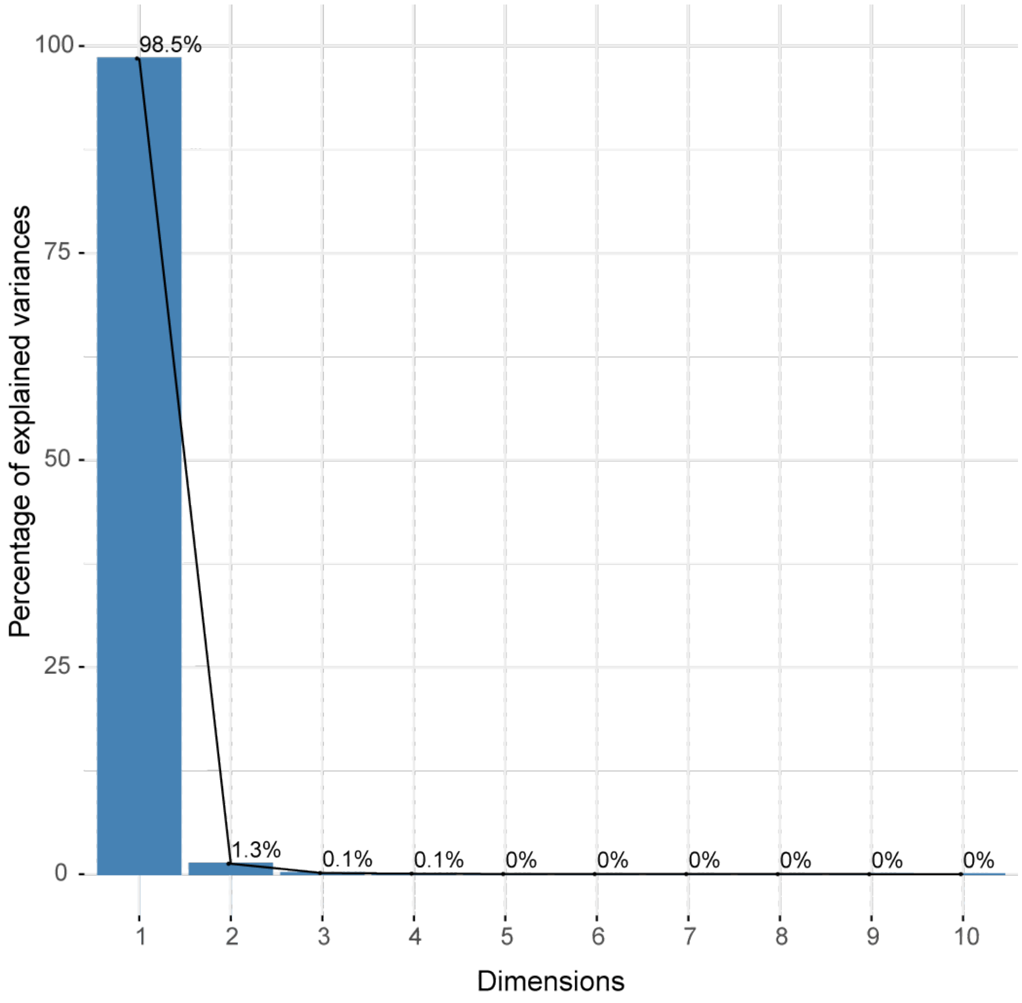


**Supplementary Figure 4 The effectiveness of each dimension in principal component analysis.** Only the first dimension (G3118T) could explain more than 98% variations.

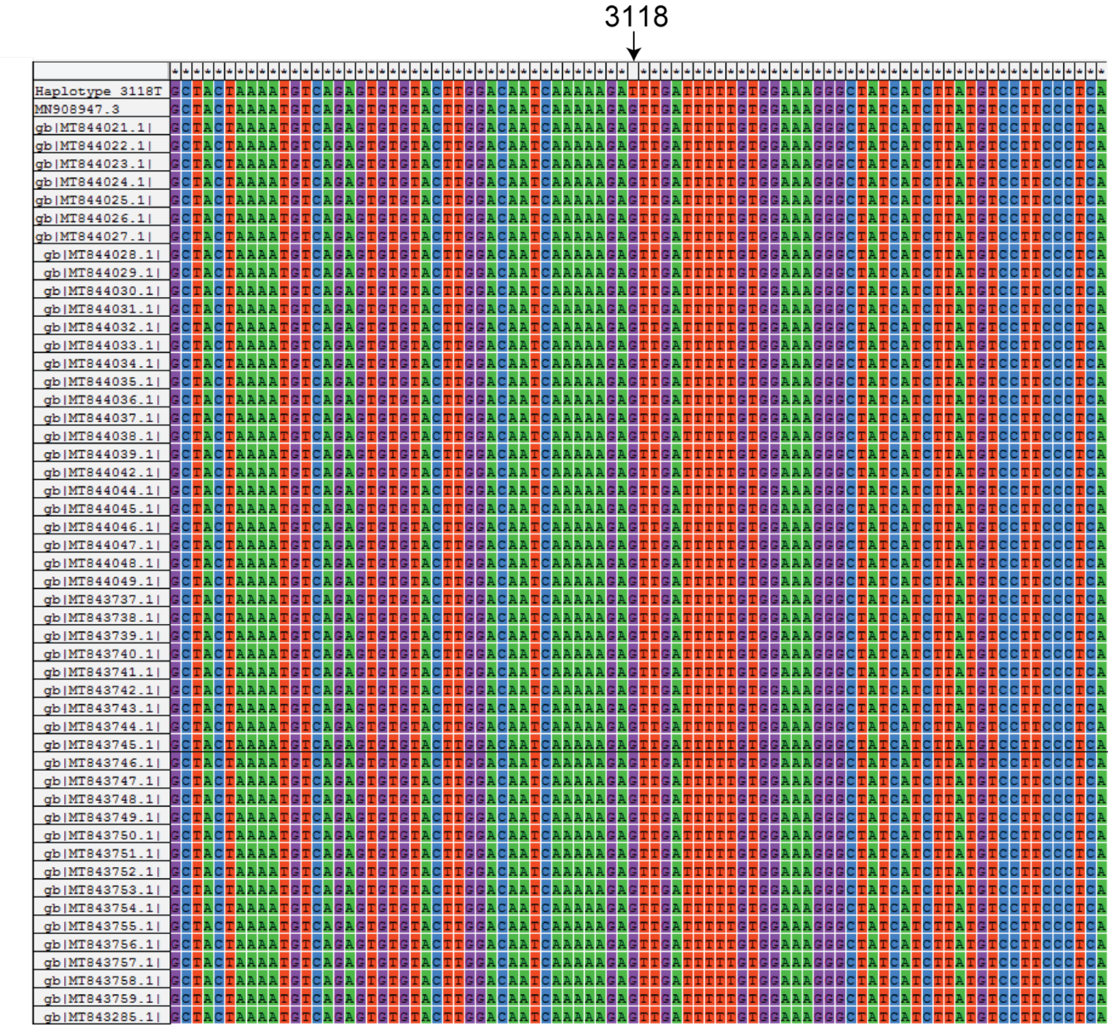


**Supplementary Figure 5 Global aligning between the haplotype 3118T and published SARS-Cov-2 genomes from GenBank.** We listed top 50 best alignment results, all SARS-Cov-2 genomes came from NCBI (Aug, 2020) except for dominant genotype T3118. The results showed that T3118 genotype was a novel dominant mutation.

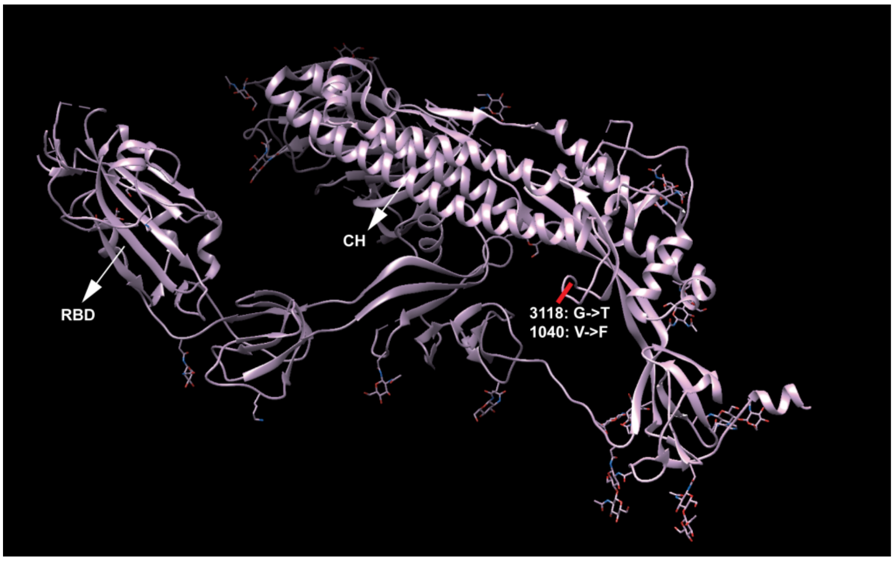


**Supplementary Figure 6 The position of three stable transmission SNVs in three-dimensional ribbon model of spike protein.** The changed amino acids are highlighted by red line and some potential functional domains are marked with arrow. The novel dominant genotype of T3118 was located in the turn site between CH and CD subdomains. Figure A, B, and C represent each of three variations with a clear vision, and figure C showed all three variations at the same time.

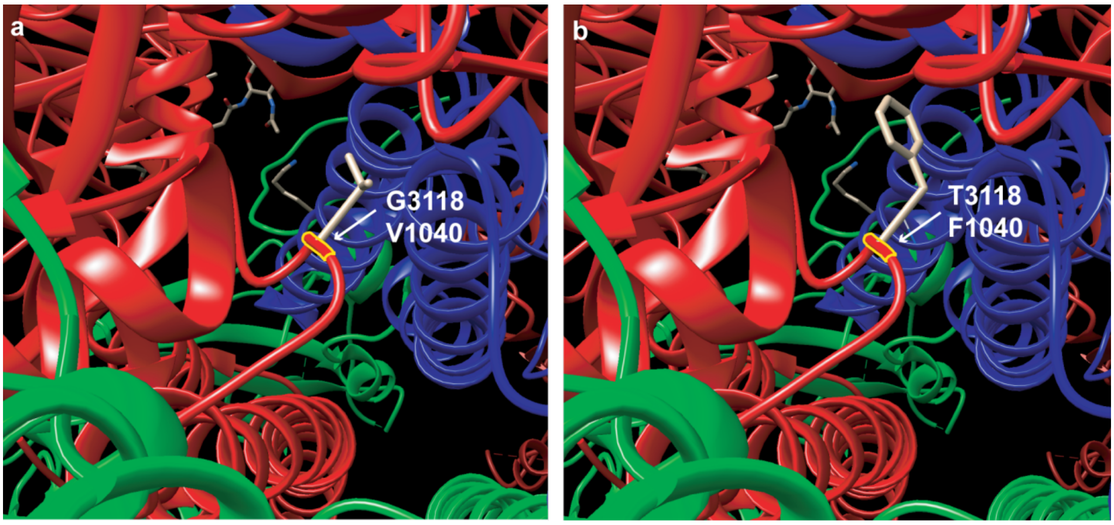


**Supplementary Figure 7 Amino acid variations of the G3118GT.** Part A represented for the genotype of SG dominant haplotype (also the early ancestor strain), and part B represented for the novel genotype of FG dominant haplotype. The location of variation site was highlighted by yellow color, the changed angle of R-group was simulated by Chimera with the Posterior probability=0.30.
